# An Open-Source Platform for Head-Fixed Operant and Consummatory Behavior

**DOI:** 10.1101/2023.01.13.523828

**Authors:** Adam Gordon-Fennell, Joumana M. Barbakh, MacKenzie Utley, Shreya Singh, Paula Bazzino, Raajaram Gowrishankar, Michael R. Bruchas, Mitchell F. Roitman, Garret D. Stuber

## Abstract

Head-fixed behavioral experiments in rodents permit unparalleled experimental control, precise measurement of behavior, and concurrent modulation and measurement of neural activity. Here we present OHRBETS (Open-Source Head-fixed Rodent Behavioral Experimental Training System; pronounced ‘Orbitz’), a low-cost, open-source ecosystem of hardware and software to flexibly pursue the neural basis of a variety of motivated behaviors. Head-fixed mice tested with OHRBETS displayed operant conditioning for caloric reward that replicates core behavioral phenotypes observed during freely moving conditions. OHRBETS also permits for optogenetic intracranial self-stimulation under positive or negative operant conditioning procedures and real-time place preference behavior, like that observed in freely moving assays. In a multi-spout brief-access consumption task, mice displayed licking as a function of concentration of sucrose, quinine, and sodium chloride, with licking modulated by homeostatic or circadian influences. Finally, to highlight the functionality of OHRBETS, we measured mesolimbic dopamine signals during the multi-spout brief-access task that display strong correlations with relative solution value and magnitude of consumption. All designs, programs, and instructions are provided freely online. This customizable ecosystem enables replicable operant and consummatory behaviors and can be incorporated with methods to perturb and record neural dynamics *in vivo*.

**Impact Statement:** A customizable open-source hardware and software ecosystem for conducting diverse head-fixed behavioral experiments in mice.

## Introduction

Studying mouse behavior under head-fixed conditions offer many distinct advantages over freely moving conditions. Head-fixation offers high degrees of behavioral control that enables consistent delivery of stimuli to the animal, precise measurement of behavior, and isolation of subcomponents of behavior (Bjerre & Palmer, 2020). Holding the mouse stable permits a wide range of behavioral experiments that have features that are challenging or impossible to conduct reliably in the freely moving condition including the delivery of somatosensory stimuli to select locations, temporally precise odor delivery (Han et al., 2018), presentation of visual stimuli to fixed parts of the visual field (Krauzlis et al., 2020; The International Brain Laboratory et al., 2021), temperature manipulations (Jung et al., 2022), and high-resolution video recording of facial expression or paw movement (Dolensek et al., 2020; Mathis et al., 2018). Eliminating or controlling physical approach behaviors also allows for isolation of both appetitive and consummatory behaviors and related neuronal dynamics. By removing turning associated with locomotion, head-fixed behavioral approaches offer enhanced compatibility with neuroscience approaches that require tethers including optogenetics and fiber-photometry. Furthermore, head-fixation is also compatible with tools for measuring and manipulating neuronal activity at the single cell level, including two-photon calcium imaging and holographic optogenetics.

Motivated behaviors are essential for survival and can be disrupted in brain circuits, leading to various diseases such as addiction and obesity. (Kenny, 2011; Rossi and Stuber, 2018; Volkow et al., 2017). Motivation in animal models is often assessed and quantified using multiple tasks that attempt to isolate distinct behavioral components such as appetitive and consummatory behaviors that can be the product of independent or overlapping brain circuits (Panksepp, 1982; Robinson and Berridge, 1993). To determine the role of brain circuits in distinct components of behavior, behavioral models with a high degree of experimental control and reproducibility are paramount as they can isolate components of behavior and limit variability across labs, subjects, and trials. There are a variety of approaches in freely moving rodents that model individual components of motivated behavior. Motivation is often modeled using operant responding on levers or nose pokes to earn a caloric reward or intracranial brain stimulation. A highly controlled version of operant responding includes retractable levers and retractable lick-spouts to limit access of both operant and consummatory responses, respectively. In contrast, consummatory behaviors require measuring the volumetric amount of appetitive or aversive solutions. A particularly useful model is the brief-access task, which consists of trial-based presentations of one of multiple solutions, enabling recording of behavioral and neuronal responses to gradations of both rewarding and aversive solutions within a single session (Boughter et al., 2002; Davis, 1973; Smith, 2001). Despite the widespread use of these procedures in freely moving animals, there has been limited adaptation of these tasks for head-fixed rodents despite advantages.

Here, we present OHRBETS, a low-cost, open-source ecosystem of hardware and software for quantifying both operant and consummatory behavior in head-fixed mice. OHRBETS features the ability to precisely limit operant and consummatory behaviors during operant conditioning, replicating the retractable levers and spout aspects of the freely moving condition. OHRBETS has a multi-spout design that allows multiple solutions to be presented independently in a single behavioral session, enabling various behavioral experiments like probabilistic reinforcement tasks and choice behavior. The platform is also flexible and includes connectivity for additional customizable components. OHRBETS consists largely of 3D printed and low-cost components that reduce the total cost per system and maximizes reproducibility. Multiple research groups have developed models for head-fixed operant behavior with a variety of operant responses (Bloem et al., 2022; Cui et al., 2017; Guo et al., 2014; Stephenson-Jones et al., 2020; Vollmer et al., 2022, 2021), but many of these systems are built for a single experimental procedures with minimal publicly available resources needed for consistent replication. To assist with modification and reproduction of our system, we have created a GitHub repository (https://github.com/agordonfennell/OHRBETS) that contains 3D models (also available through TinkerCad), assembly instructions, wiring diagrams, behavioral programs, and scripts for analysis. The OHRBETS ecosystem will allow any investigator to harness the strength of head-fixed approaches to study the neurobiological underpinnings of motivation and related disease states while maintaining many crucial behavioral phenotypes established in freely moving animals.

## Results

### OHRBETS Overview

We developed OHRBETS, a low-cost, open-source system for head-fixed behaviors in mice (**Figure 1A-E**, **Figure 1- figure supplement 1**). Our system consists of custom 3D printed and inexpensive, commercially available components bringing the total cost to around $600 for the operant- only version and around $1,000 for the operant + multi-spout version. For head-fixation, mice are implanted with a metal head-ring and are easily and quickly secured on the head-fixed system for daily behavioral sessions (**Figure 1A-B**). To deliver solutions, including sucrose, we use gravity fed tubing attached to a stainless-steel lick spout that is gated by a solenoid. The position of the lick spout is controlled using a custom 3D printed micropositioner (Backyard Brains 2013; Hietanen et al., 2018), and licks are detected using a capacitive touch sensing. To limit access to consumption, paralleling a retractable lick spout from the widely used freely moving operant assay, we used a linear actuator (adapted from (Buehler, 2016a)) that is controlled using a 5V micro servo for extending and retracting the spout (**Figure 1D**). A 43.2 mm diameter wheel (Lego, 86652c01) coupled to a rotary encoder is mounted underneath the mouse such that their forepaws’ deflections left or right can serve as the operant response (International Brain Laboratory 2021). To limit access to operant responding, paralleling retractable levers used in the freely moving operant assay, we developed a wheel brake controlled via an additional micro servo (**Figure 1E**). All behavioral components are controlled by an Arduino Mega and the timing of events are relayed via serial communication and recorded using a Python program (**Figure 1C**). Our system is inexpensive and easily assembled following instructions freely available through our GitHub repository (https://github.com/agordonfennell/open_stage).

**Figure 1:**
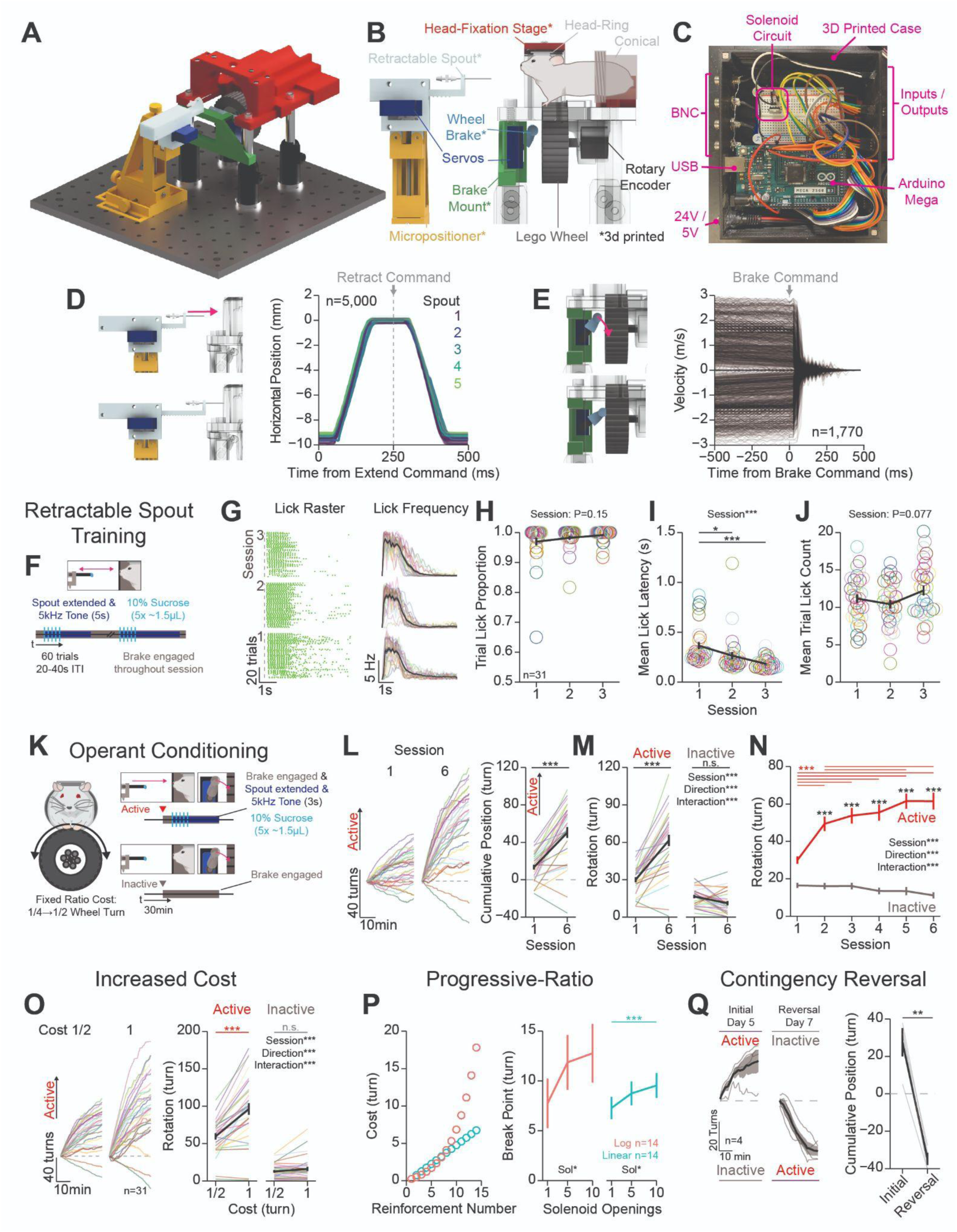
Mice Rapidly Learn Head-Fixed Operant Conditioning for Sucrose and Display Operant Behaviors Established in Freely moving Experiments: (A-E) System overview. (A) 3D rendering of our open-source, low-cost, head-fixed system (Open-Source). B) Cartoon depicting the critical components of our system (* indicates 3D printed components). C) Image of the Arduino based microprocessor and custom enclosure used for controlling hardware and recording events. D) Validation of our 3D printed retractable spout powered by a low-cost micro servo. left: 3D rendering of the linear travel of the spout; right: horizontal position of the spout tip determined using DeepLabCut over time during 1,000 extension/retractions with 5 unique retractable spout units. E) Validation of our 3D printed wheel brake powered by a low-cost micro servo. 3D rendering of the rotational travel of the wheel brake (left); binned rotational velocity of the wheel produced by manual rotation before and after the brake is engaged (right). F) Cartoon depicting the task design for retractable spout training. G) Licking behavior throughout retractable spout training; lick raster for a representative mouse with each lick represented as a tick (left); mean binned frequency of licks (right). (H-J) Summary of behavior throughout retractable spout training: proportion of trials with at least 1 lick (H); Mean latency from spout extension command to first lick on trials with a lick (I); mean number of licks within each 5 s access period (J). K) Cartoon depicting the task design for operant conditioning. L) Cumulative position of the wheel throughout the session (left) and at the conclusion of the session (right) on the first and sixth session of training (Positive direction indicate rotation in the active direction; session 1 vs session 6 *t*-test***). (M-N) Total rotation of the wheel throughout a session broken down based on direction on the first and sixth session of training (M) and across training sessions (N). O) Cumulative position of the wheel throughout the last session (left), and the mean total rotation of the wheel in the last 3 sessions of fixed-ratio 1/2 turn and 1 turn. P) Progressive-ratio schedule of reinforcement (left) and break points across different reward magnitudes set by the number of solenoid openings (One-Way RM ANOVA*). Q) Cumulative position of the wheel throughout the session (left) and at the conclusion of the session (right) on the last session of initial training and reversal training (*t*-test; initial vs. reversal: *t*-test**). (*Unless otherwise noted, effects listed on plots indicate statistical significance for Two-Way RM ANOVA effects; Multi color lines and rings depict individual mice; Black lines depict mean across mice; Black asterisks above horizontal bars in (N) and (P) indicate significant differences in active rotation across sessions, while black asterisks above means indicate significant differences between active and inactive rotation within a session; see stats table for details*).

To characterize the effectiveness of our retractable spout and wheel brake, we conducted experiments to determine the timing and reliability of the hardware. We measured the linear travel of 5 sets of retractable spouts using high speed video recording (200 fps) during 1000, 1 cm spout extensions/retractions and determined the position of the spout using DeepLabCut (Mathis et al., 2018) (**Figure 1D**, **Figure 1- figure supplement 1B**). We found that the retractable spout follows a consistent and reliable pattern with >98% of extensions reaching a terminal position within 0.3 mm of each other in under 180ms of the extension command (**Figure 1A**, **Figure 1- figure supplement 1B A-E**). We measured the braking ability of 4 sets of wheel brakes by manually rotating the wheel at different rates in both directions and then programmatically engaging the brake (**Figure 1E**, **Figure 1- figure supplement 1G**). The wheel brake rapidly stopped wheel rotation in 100% of trials, even with manual velocities that exceed that which a mouse can produce (**Figure 1E**). Furthermore, we analyzed the effectiveness of the brake to stop wheel rotation in data obtained during operant conditioning experiments and found that most mouse-generated rotations ceased in under 250ms (**Figure 1- figure supplement 1**). Together, these results indicate OHRBETS produces reliable spout extension/retraction and wheel braking using inexpensive micro servos and 3D printed components, and therefore will effectively limit access to consummatory and operant responses during behavioral experiments.

### OHRBETS trained mice show multiple established characteristics of operant behavior observed in freely moving animals

We developed a training procedure that permits measuring operant conditioning in head-fixed mice, and we conducted a series of experiments to determine if operant behavior conducted with OHRBETS reproduces behavior seen in freely moving rodents (Kliner et al., 1988; Reilly, 1999; Winger and Woods, 1985). We trained head-fixed, water-restricted mice to perform operant conditioning in 3 stages: 1) free-access lick training, 2) retractable spout training, and 3) operant conditioning (*Methods*). To measure the reproducibility of OHRBETS, all experiments were conducted using 4 independent Operant-Stage assemblies (referred to as box ID, data shown in supplements).

We trained mice on a single session of free-access lick training to facilitate licking from the spout and reduce stress associated with head-fixation (**Figure 1- figure supplement 2**). Free-access lick training consisted of a 10 min session where each lick immediately triggered a delivery of ∼1.5 µL of 10% sucrose which approximates free-access consumption from a standard lick spout (**Figure 1- figure supplement 2A**). During training, 100% of mice licked for sucrose throughout the session (**Figure 1- figure supplement 2B**). Like the standard freely moving free-access assay (Johnson, 2018; Spector et al., 1998), OHRBETS trained mice licked in discrete licking bouts (**Figure 1- figure supplement 2C-D**). The total number of licks as well as the licking microstructure, including licks per bout and bout duration, were consistent across sex, cohort, and box ID (*t*-test or One-Way RM ANOVA n.s.; **Figure 1- figure supplement 2E-P**), as well as across freely moving and head-fixed conditions (*t*-test n.s.; **Figure 1- figure supplement 2Q-V**).

Next, mice completed 3 sessions of retractable lick spout training - building the association between spout extension and the availability of reward to enhance the learning rate in subsequent operant conditioning (Steinhauer et al., 1976) (**Figure 1F-J**, **Figure 1- figure supplement 3**). Each session consisted of 60 trials of spout extension, delivery of 5 pulses of 10% sucrose (∼1.5 µL/pulse, 200ms inter-pulse interval), and a 5 s access period for liquid to be consumed during which a 5 kHz tone was presented. Mice licked to consume sucrose delivered on most trials with a short latency between spout extension and licking throughout each session (**Figure 1G**, **Figure 1- figure supplement 3A-E**). By the third session of training, 31 out of 31 mice licked during 90% of trials (**Figure 1H**). Mice demonstrated a learned association between spout extension and a simultaneous auditory tone with the availability of sucrose, as they reduced their latency from spout extension to first lick across the 3 sessions of training (Two-Way RM ANOVA: Session***, **Figure 1I**). No changes in the proportion of trials with a lick or the number of licks per trial over sessions were observed (One-Way RM ANOVA: Session n.s.; **Figure 1I, J**). Female mice displayed a higher lick latency in response to spout extension compared to males on the first session of training (Two-Way RM ANOVA: Sex**, Sex x Session**; Session 1 HSD***), but the proportion of trials with a lick and the number of licks per trial was not statistically different between males and females (Two-Way RM ANOVA: Sex n.s., Sex x Session n.s.; **Figure 1- figure supplement 3E, H, K**). There were no differences in behavior across cohorts or behavioral systems (Two-Way RM ANOVA: main effects and interactions n.s..; **Figure 1- figure supplement 3F, I, G, J, M**), aside from a significant interaction between cohort and session for the mean trial lick count (Two-Way RM ANOVA: Cohort x Session**; **Figure 1- figure supplement 3L**). These data indicate that mice rapidly learn to lick for sucrose during discrete windows of access.

After free-access lick training and retractable spout training, water-restricted mice were operantly conditioned for sucrose (**Figure 1K-N**, **Figure 1- figure supplement 4**). Operant conditioning consisted of 6 sessions of responding for 10% sucrose under fixed-ratio schedule (1/4 rotation for session 1; 1/2 rotation for sessions 2-6; **Figure 1K**; **Figure 1- Figure Supplement 6**; *Methods*). To assess if mice learned the operant requirement, we examined whether mice increased responding in the active direction over sessions and exhibited a response bias for the active over the inactive response (Heyser et al., 2000). We found that mice learned to turn the wheel to obtain 10% sucrose in as little as 1 session, as 25/31 mice showed greater rotation in the active direction compared to the inactive direction (**Figure 1L, session 1**). By the 6^th^ session of operant conditioning, 29/31 mice showed an increase in net rotation in the active direction (*t*-test***; **Figure 1L**, data from all sessions shown in **Figure 1- figure supplement 4A**), that was the product of increased rotation in the active direction and no change in rotation in the inactive direction (Two-Way RM ANOVA: Session x Direction***; **Figure 1M**). As a group, mice showed significantly more rotation in the active direction compared to the inactive direction starting at the second session (Two-Way RM ANOVA: Session x Direction***; Session 2-6 Active vs Inactive: HSD***; **Figure 1N**). Mice that were tested in each of the 4 boxes showed similar inter-lick intervals, trial lick counts, and latency to lick (**Figure 1- figure supplement 4B-D**). When analyzing behavioral data based on sex, cohort, and box ID, we found only minor differences in behavior (**Figure 1- figure supplement 4E-Y**). Notably, we found that over the course of training sessions, female mice exhibit a reduced total active rotation (Two-Way RM ANOVA: Sex*; **Figure 1- figure supplement 4H**), reduced total lick count (Two- Way RM ANOVA: Sex*; **Figure 1- figure supplement 4Q**), and reduced bias for rotation in the active direction (Two-Way RM ANOVA: Sex*; **Figure 1- figure supplement 4T**). These data indicate that mice rapidly exhibit operant responding for sucrose using OHRBETS, and this behavior is consistent across training history and behavioral setup with only minor differences observed a between males and females.

Next, we determined if OHRBETS could reproduce other behaviors that have been established in freely moving rodents, including increased active responding following increased cost of reward (Kliner et al., 1988; Winger and Woods, 1985) (**Figure 1O**), progressive-ratio responding with a fixed reward magnitude (Reilly, 1999; Sclafani and Ackroff, 2003; Winger and Woods, 1985) (**Figure 1P**), and reversal learning (Forgays and Levin, 1959; Heyser et al., 2000; Klanker et al., 2015) (**Figure 1Q**). To measure the relationship between cost and active response rate, after completing 1 session with a fixed-ratio of 1/4 turn and 5 sessions of a fixed-ratio of 1/2 turn (**Figure 1L-N**), we increased the fixed-ratio to 1 turn and measured operant responding for 4 sessions. As observed in freely moving rodents (**Figure 1- figure supplement 5B**), when we increased the cost of reward, mice significantly increased responding in the active direction but not the inactive direction (Two-Way RM ANOVA: Cost x Direction***; **Figure 1O**, all sessions shown in **Figure 1- figure supplement 5A**) indicating that they show flexible response rates as a function of reward cost (Kliner et al., 1988; Winger and Woods, 1985). Next, to measure the motivation to seek different reward magnitudes, we tested mice over multiple sessions of progressive ratio responding for sucrose of varying volumes (1, 5, 10 deliveries of ∼1.5 µL of 10% sucrose, counterbalanced order). During progressive-ratio sessions, mice were tested with a linear or logarithmic reinforcement schedule, where the cost for each subsequent reinforcer was higher than the last (**Figure 1P** **left**, *Methods*). Under both schedules, mice responded for rewards during progressive ratio and displayed increased breakpoints for greater reward magnitude (Reilly, 1999; Sclafani and Ackroff, 2003; Winger and Woods, 1985) (One-Way RM ANOVA: Number of Solenoid Openings*; **Figure 1P** **right**). To determine if mice can learn reversals in response contingency, we trained a naïve group of mice to perform operant responding with an initial rotational direction contingency for 5 sessions and then switched the contingency and allowed mice to re-learn over 7 sessions. We found that mice displayed reversal learning, as they reversed the terminal cumulative position (initially active - initially inactive) following contingency reversal and training over 7 sessions (Heyser et al., 2000; Klanker et al., 2015)(*t*- test**; **Figure 1Q**). Finally, to directly compare behavior during head-fixed and freely moving versions of operant conditioning, we examined behavioral responding in the two tasks within the same mice (**Figure figure supplement 5A, b**). We found that mice showed similar changes in response vigor with increased cost of reward (**Figure 1- figure supplement 5A-C**) and similar pattern of reduction in responding over the course of a session after the first 10 min (**Figure 1- figure supplement 5D**) but earned more liquid in the freely moving version of the task (*t*-test***; **Figure 1- figure supplement 5E**). Together, these data indicate that mice display flexible operant behavior in our head-fixed system that is sensitive to the cost of reward, the magnitude of reward, and reward contingency, and produces behavior in a parallel manner to freely moving operant conditioning.

### OHRBETS trained mice exhibit positive and negative operant conditioning during optogenetic stimulation of LHA GABAergic and glutamatergic neurons

After establishing that mice display operant responding for caloric rewards, we determined if they would perform operant responding to obtain or avoid optogenetic stimulation of brain circuits that have been previously established to be rewarding or aversive in freely moving rodents using the OHRBETS system (Chen et al., 2020; Jennings et al., 2015, 2013; Rossi et al., 2019). Optogenetic stimulation allows for temporally precise manipulations of genetically- and spatially- defined neuronal circuits enabling greater consistency of unconditioned stimuli delivery across a multitude of experimental conditions. We used optogenetic stimulation of LHA GABAergic neurons (LHA^GABA^) as a appetitive unconditioned stimulus because activation of these neurons produces positive reinforcement (Jennings et al., 2015), and optogenetic stimulation of LHA Glutamatergic neurons (LHA^Glut^) as an aversive unconditioned stimulus because activation of these neurons is aversive (Chen et al., 2020; Rossi et al., 2019). To selectively manipulate LHA^GABA^ and LHA^Glut^ neurons, we expressed cre-dependent channelrhodopsin-2 (ChR2) or cre-dependent mCherry in the LHA of *Slc32a1*^Cre^ (*Vgat*-cre) or *Slc17a7*^Cre^ (*Vglut2*-cre) mice (Vong et al., 2011) and implanted bilateral optic fibers with a head-ring to facilitate head-fixation (**Figure 2A**, **Figure 2- figure supplement 1A**, *Methods*). Following incubation, mice were tested using freely moving and OHRBETS intracranial self-stimulation (ICSS) in counterbalanced order.

**Figure 2:**
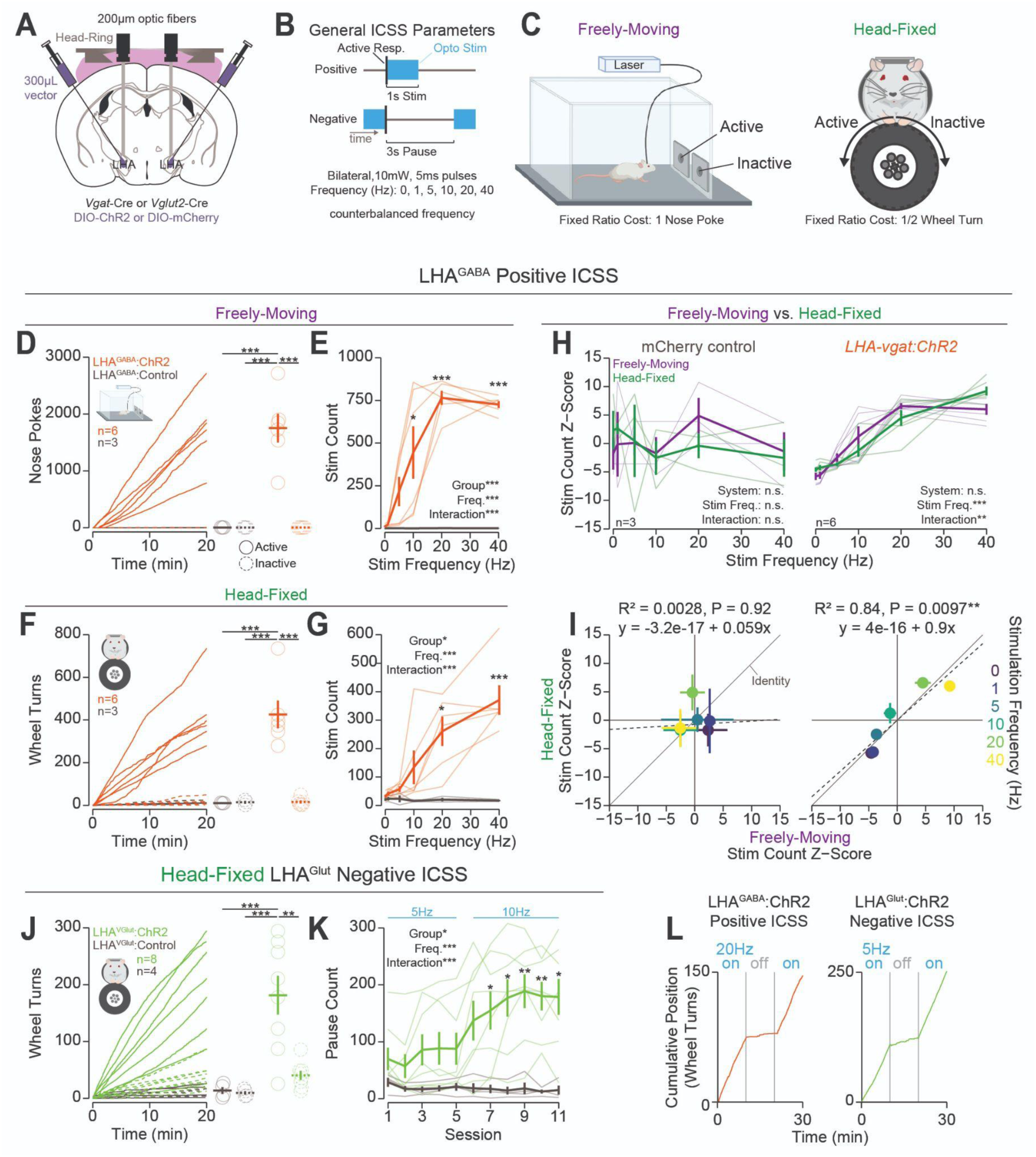
Head-fixed Operant Conditioning to Obtain Stimulation of LHA^GABA^ Neurons or Avoid Stimulation of LHA^Glut^ Neurons: A) Approach, placements depicted in Figure 2- figure supplement 1A. B) Diagram of the experimental approach for positive reinforcement in LHA^GABA^ mice and negative reinforcement in LHA^Glut^ mice. C) Cartoon depicting the freely moving (left) and head-fixed (right) versions of the operant task. D) Cumulative (left) and total (right) nose pokes under positive reinforcement for 40 Hz stimulation in LHA^GABA^:ChR2 (red) and LHA^GABA^:Control (grey) mice. E) Total number of stimulations earned under positive reinforcement for multiple stimulation frequencies (1 frequency/session). (F, G) same as (D, E) except during the head-fixed version of the task. H) Comparisons of the *z*-score of the total number of stimulations across frequencies in freely moving (purple) and head-fixed (green). *Z*-scores were calculated for each mouse x system independently. No significant *post hoc* differences when comparing systems at the same stimulation frequency. I) Correlation of the *z*-score of the total number of stimulations in the freely moving and head-fixed version of the task. J) Cumulative rotation over a session under negative reinforcement for 5 Hz and 10 Hz stimulation in LHA^Glut^:ChR2 (lime green) and LHA^Glut^:Control (grey) mice. K) Total pause count across all training sessions (frequency schedule indicated with blue text above the plot; *asterisks above means indicate significant differences determined by Bonferroni adjusted t-test*). L) Cumulative position over a 30 min session with the laser turned off from 10-20 min. (*Unless otherwise noted*, *Effects listed on plots indicate statistical significance for Two-Way RM ANOVA effects; Faded lines and rings depict individual mice; asterisks above means indicate significant differences determined by HSD between stim count at a corresponding stim frequency or pause count at a corresponding session; asterisks above horizontal lines indicate significant difference determined by HSD between means indicated by line; see stats table for details*).

Mice displayed high levels of active responses to obtain optogenetic stimulation of LHA^GABA^ neurons that was consistent across freely moving and head-fixed procedures (**Figure 2D-I**, **Figure 2- figure supplement 1B-G**, **Figure 2- Figure Supplement 2**). We first trained mice to nose poke (fixed- ratio 1 poke) or turn a wheel (fixed-ratio 1/2 turn) to obtain optogenetic stimulation (1 s, 20 Hz, 5 ms pulse duration) of LHA^GABA^ cells over 4-5 sessions (**Figure 2B, C**; training data shown in **Figure 2- figure supplement 1B-G**). Next, we measured operant responses for different stimulation frequencies by running mice through 5 sessions of ICSS with one of 5 stimulation frequencies (1, 5, 10, 20, 40 Hz) in counterbalanced order. On the last 20 Hz self-stimulation training session, *Vgat*-cre mice expressing ChR2 in the LHA (LHA^GABA^:ChR2) displayed high levels of operant responding for the active hole or active direction and displayed strong discrimination between active and inactive responses; *Vgat*-cre mice expressing the mCherry control construct in the LHA (LHA^GABA^:Control) displayed little to no responding and did not discriminate between responses (Two-Way RM ANOVA: Response ID x Group** for both systems; **Figure 2D, F**). In both the freely moving and head-fixed conditions, LHA^GABA^:ChR2 mice displayed greater active response rates for higher stimulation frequencies (Two-Way RM ANOVA: Group x Stimulation Frequency*** for both systems; **Figure 2E, G, H**) that were positively correlated across the two versions of the task (Pearson’s Product Moment**; **Figure 2I**). On the contrary, LHA^GABA^:Control mice displayed no change in responding to changes in frequency and no correlation across the two procedures. Similar to freely moving ICSS (Stuber et al., 2011; Witten et al., 2011), mice trained with positive reinforcement rapidly ceased responding once optogenetic stimulation was withheld and then resumed responding once optogenetic stimulation was reintroduced (**Figure 2L, left**). These data indicate that OHRBETS can robustly elicit motivated behaviors to obtain rewarding optogenetic stimulation in a similar manner to freely moving rodent behavioral paradigms.

Mice displayed high levels of responding to avoid optogenetic stimulation of LHA^Glut^ neurons under negative reinforcement during the head-fixed procedure but not the freely moving procedure. To elicit negative reinforcement (responses to cease an aversive stimulus) in the head-fixed procedure, we trained mice to turn a wheel to earn a 3 s pause of continuous stimulation of LHA^Glut^ neurons at 5 Hz for sessions 1 - 5 and 10 Hz for session 6 - 11 (**Figure 2b**). Following training, *Vglut2*-cre mice with expression of ChR2 in the LHA (LHA^Glut^:ChR2) displayed high levels of responding in the active direction and strong discrimination between the active and inactive directions, while *Vglut2*-cre mice with expression of mCherry control construct in the LHA (LHA^Glut^:Control) displayed little to no responding and no discrimination (Two-Way RM ANOVA: Direction x Group*; **Figure 2J**, **Figure 2- Figure Supplement 2**). Over the course of training, LHA^Glut^:ChR2 mice, but not LHA^Glut^:Control mice, increased the number of pauses earned (Two-Way RM ANOVA: Stimulation Frequency x Group***; **Figure 2K**). Compared to LHA^Glut^:Control, LHA^Glut^:ChR2 mice showed substantially higher active rotation as well as a moderately higher inactive rotation (**Figure 2- figure supplement 1K, M**). LHA^Glut^:ChR2 mice acquired negative reinforcement behavior at a reduced rate compared to positive reinforcement (7 sessions to acquisition of negative reinforcement vs 1 session for positive reinforcement; **Figure 2K**, **Figure 2- figure supplement 1F, L**). Like positive reinforcement, LHA^Glut^:ChR2 mice that were trained on negative reinforcement rapidly ceased responding when the optogenetic stimulation was removed and resumed responding when optogenetic stimulation was reintroduced (**Figure 2L, right**). To compare behavior in head-fixed to freely moving procedures, we trained the same mice under negative reinforcement in a freely moving procedure. We found that, compared to LHA^Glut^:Control mice, LHA^Glut^:ChR2 mice displayed suppressed amounts of active-responding, number of pauses earned, and inactive responding during the freely moving condition (**Figure 2- figure supplement H-J**). The discrepancy between acquisition of negative reinforcement in the head-fixed assay versus the freely moving assay could be attributed to the reduced range of actions mice can make in the head-fixed assay. These results indicate that OHRBETS can elicit responding under negative reinforcement using a simple stimulation procedure that is incapable of producing responding in traditional freely moving conditions.

### Head-fixed mice express real-time place preference and avoidance behaviors

We designed and tested a procedure analogous to real-time place testing (RTPT) (Britt et al., 2012; Kravitz et al., 2012; Stamatakis and Stuber, 2012; Tye and Deisseroth, 2012) in head-fixed mice (**Figure 3**). RTPT is extensively used to measure the appetitive or aversive characteristics of neuronal manipulations. With the same mice utilized for operant conditioning (*Methods*), we used stimulation of LHA^GABA^ neurons as a positive unconditioned stimulus and stimulation of LHA^Glut^ neurons as a negative unconditioned stimulus because these two populations have been previously shown to drive real-time place preference (RTPP) and real-time place avoidance (RTPA), respectively (Jennings et al., 2015; Nieh et al., 2016; Rossi et al., 2019) (**Figure 3A**, for fiber placement see **Figure 3- figure supplement 1A**). *Vgat*-cre and *Vglut2*-cre mice expressing mCherry were pooled after observing no statistical differences in behavior between the two genotypes. In the standard version of the task, freely moving mice traverse a two-chamber arena in which they receive optogenetic stimulation when the mouse is located in one of the two chambers (**Figure 3B, top**). Using OHRBETS, the response wheel was divided into two halves relative to the starting position of the wheel, one of which was paired with optogenetic stimulation (**Figure 3B, bottom**). To enhance the mouse’s ability to determine their position on the wheel, we included a tone that indicated the mouse’s position in the two zones. The two chamber RTPT assay offers a distinct advantage for comparing behavior across different versions of the assay because throughout the entire session duration the subject is in one of two states (stimulated or not), allowing for a one-to-one comparison of the amount of time stimulated over the duration of the fixed session. For both tasks, mice were initially habituated without stimulation for 1 session and then underwent RTPT over 6 sessions with frequency and chamber/wheel-zone pairing counterbalanced (**Figure 3C**). For the head-fixed procedure, mice were initially trained without any cue indicating the wheel zone. After initial training, we paired the wheel zones with tones and found that mice exhibited more obvious RTPP/RTPA (**Figure 3- figure supplement 1D**), so in subsequent sessions these zone cues were added to the task design. Using this approach, we measured the similarity in RTPT behavior with a range of rewarding and aversive stimulation magnitudes across freely moving and head-fixed procedures.

**Figure 3:**
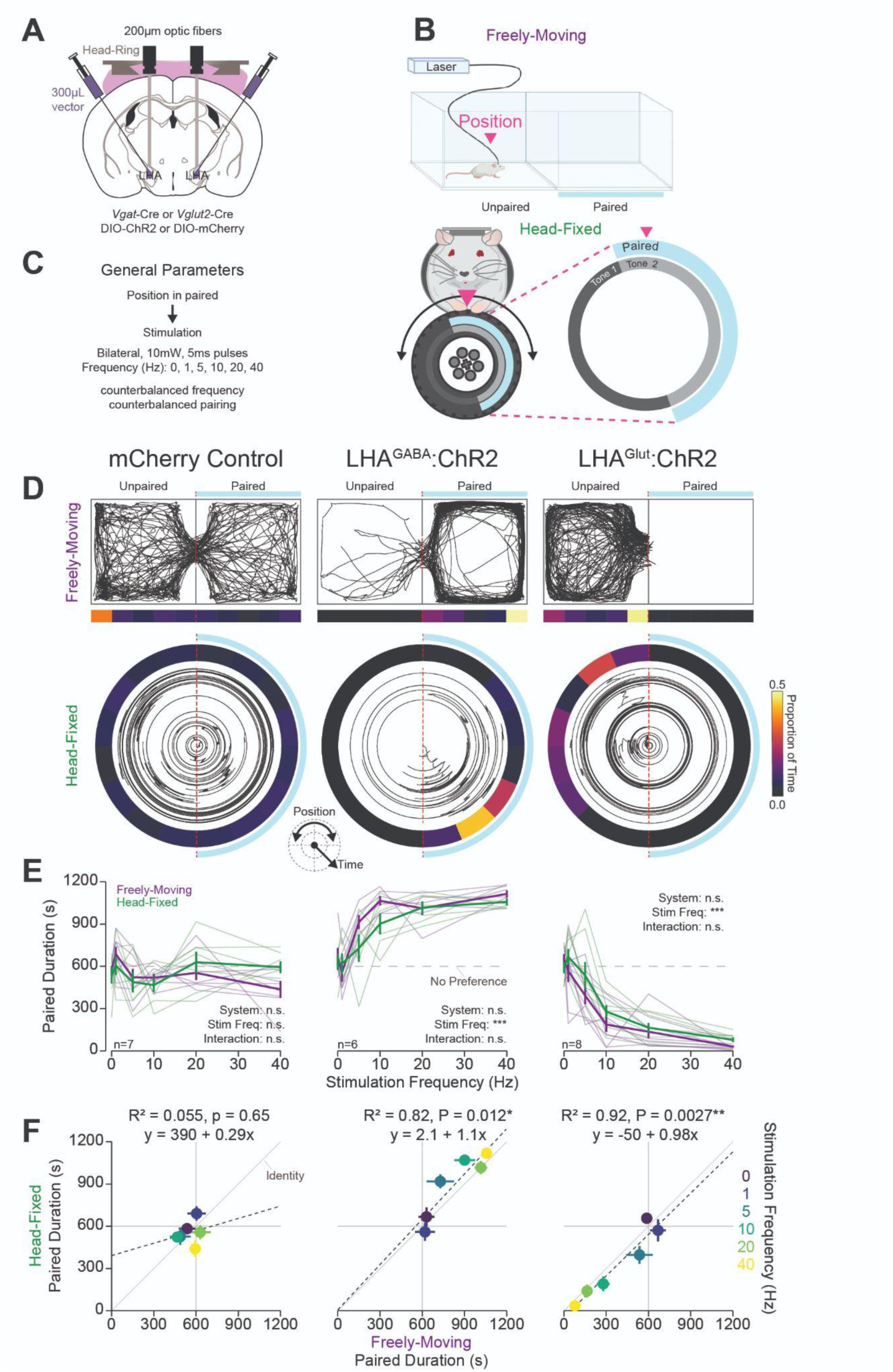
Head-fixed Real-Time Place Preference and Aversion Associated with Stimulation of LHA Subpopulations Mirrors Freely moving Behavior: A) Approach, placements depicted in Figure 2- figure supplement 1A. B) Cartoon depicting the freely moving and head-fixed versions of the operant task. In the head-fixed task, the mouse’s position was determined relative to the position of the wheel and the mouse could rotate the wheel to navigate through the paired and unpaired zones. C) Task design. (D-F) Behavior during the RTPT task; left column contains data from mCherry controls (both LHA^GABA^:Control and LHA^Glut^:Control), middle contains LHA^GABA^:ChR2, right contains LHA^Glut^:ChR2. D) Representative traces of the mouse’s position in the 2 chamber arena in freely moving RTPT (top) and the position of the wheel over time in head-fixed RTPT (bottom). The right side of the arena or wheel was paired with optogenetic stimulation as indicated by the blue bar/arc. The proportion of time in binned areas of the arena or wheel are shown in the heat maps under or surrounding the traces (color scale represents the proportion of time in each position bin). E) Amount of time spent in the paired zone during a 20 min (1200 s) session for varying frequencies; values above 600 s are indicative of preference, values below are indicative of avoidance. Colors represent the version of the task as indicated in the left column. F) Correlation between the mean time spent in the paired zone across mice (5 values) during freely moving (abscissa) and the head-fixed (ordinate) versions of RTPT at different stimulation frequencies (colors represent stimulation frequency; error bars represent SEM). (In *(E) asterisks depict Two-Way RM ANOVA effects, no HSD differences between systems were detected at corresponding stimulation frequencies; see stats table for details*).

Mice expressed similar behaviors in the RTPT assay during freely moving and head-fixed procedures (**Figure 3D-F**, **Figure 3- figure supplement 1**). Specifically, mice expressing mCherry (LHA:Control mice) did not show preference nor aversion for the stimulation paired chamber/zone across all stimulation frequencies in both the freely moving and head-fixed procedures and did not show correlations across the two assays (Two-Way RM ANOVA: Stimulation Frequency n.s.; **Figure 3D-F**). On the contrary, LHA^GABA^:ChR2 mice showed strong place preference while LHA^Glut^:ChR2 mice showed strong place aversion for the paired chamber/zone with higher stimulation frequencies compared to lower stimulation frequencies (Two-Way RM ANOVA: Stimulation Frequency*** for both groups; **Figure 3D, E**). There was no statistical difference between the amount of time in the paired chamber/zone in the freely moving and head-fixed versions of the task (Two-Way RM ANOVA: System n.s. for both groups, Stimulation Frequency x System n.s. for both groups; **Figure 3D, E**). Furthermore, the time in the paired chamber/zone was correlated across freely moving and head-fixed procedures for LHA^GABA^:ChR2 and LHA^Glut^:ChR2 mice, but not LHA:Control mice (Pearson’s Product moment: * for LHA^GABA^:ChR2, ** for LHA^Glut^:ChR2; **Figure 3F**). Examining the correlation of behavior between the two assays at different stimulation frequencies in individual mice (**Figure 3- figure supplement 1B**) revealed that, compared to LHA:Control mice, LHA^GABA^:ChR2 mice and LHA^Glut^:ChR2 mice showed a greater Pearson’s Product Moment R and regression slope estimate (**Figure 3- figure supplement 1C, left-mid**). Furthermore, LHA^Glut^:ChR2 mice showed lower regression *p*-values compared to mCherry controls while the LHA^GABA^:ChR2 mice displayed a trend towards lower *p*-values (**Figure 3- figure supplement 1C, right**). Together these results indicate that OHRBETS elicits RTPT behavior similar to freely moving procedures and provides a useful experimental approach for measuring the valence of stimuli.

### OHRBETS trained mice display consummatory behaviors dependent on the concentration of appetitive and aversive solutions

Exposure to appetitive and aversive taste solutions provides an approach to measure neuronal correlates of appetitive and aversive events in addition to operant responding. Within-session consumption of unpredictable tastants allows for measuring a range of behavioral and neuronal responses to gradations in solution valence. We adapted OHRBETS to include a retractable, radial multi- spout consisting of 5 spouts (**Figure 4A**, **Figure 4- Figure Supplement 3**). Using this system, we delivered up to 5 solutions with different concentrations in the same session with a task design adapted from the Davis Rig (Davis, 1973; Smith, 2001). Each behavioral session consisted of 100 trials with 3 s of free-access consumption separated by 5 – 10 s inter-trial intervals during which all spouts retracted (**Figure 4B**). The 5 solutions were delivered in pseudorandom order such that each solution was delivered 2 times every 10 trials. To control for modest spout effects (**Figure 4- figure supplement M- O**), we conducted the experiment counterbalanced over 5 sessions such that each spout was paired with each concentration (**Figure 4C, J, Q**). Using this approach, we measured within-session consumption of gradations in concentration of an appetitive solution (sucrose) and two aversive solutions (quinine and hypertonic sodium chloride (NaCl)).

**Figure 4:**
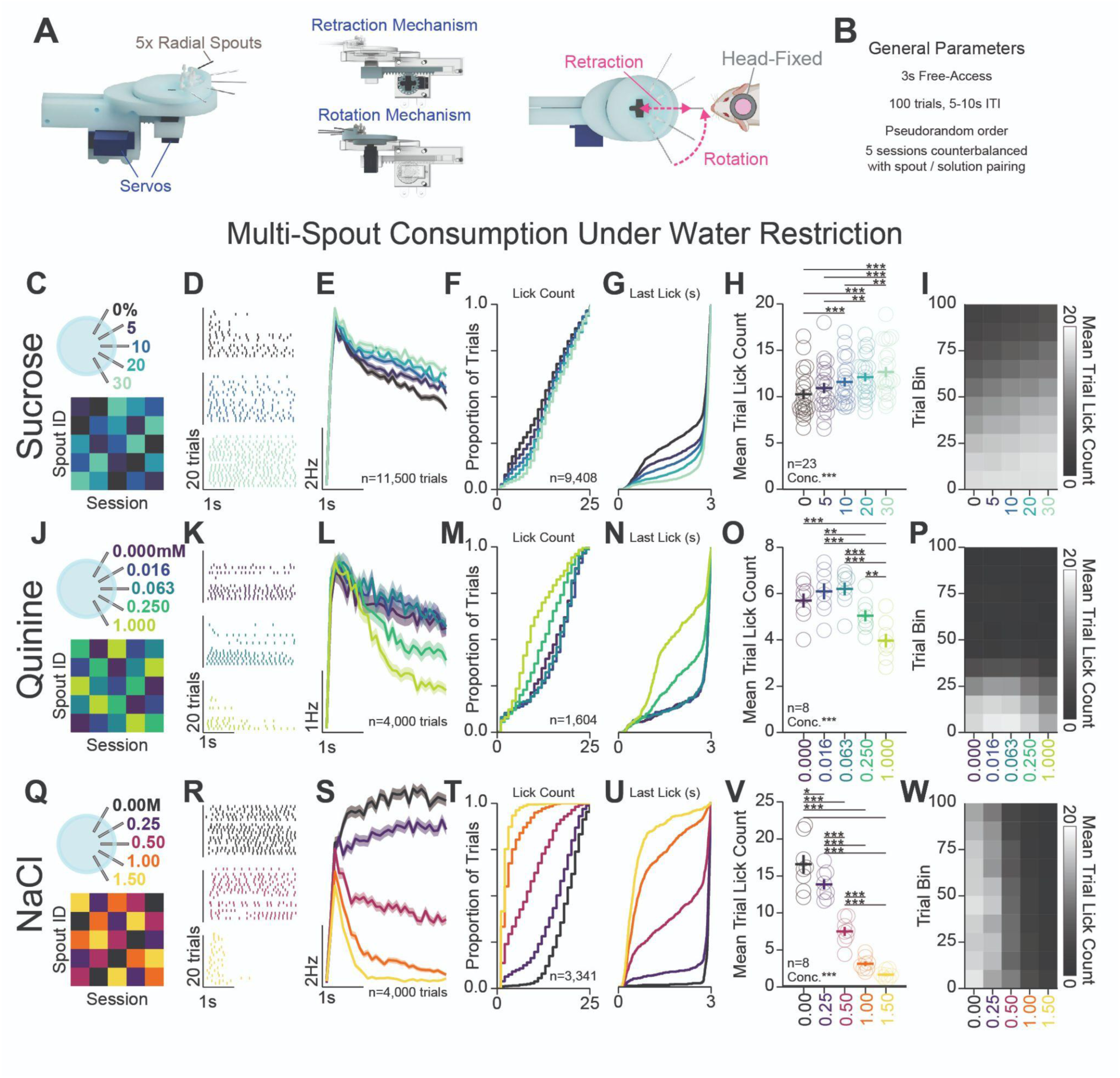
Head-Fixed Consumption of Gradients of Rewarding and Aversive Solutions During Brief Access: A) 3D rendering of the multi-spout unit that retracts and rotates to allow brief access periods to 1 of 5 lick spouts to the head-fixed mouse. B) Task design. (C-I) Multi-spout consumption of a gradient of concentrations of sucrose data. C) Procedure: mice received 5 sessions of 5x multi-spout counterbalanced to have each solution of each spout once. Colors represent concentrations of solution as defined in the label adjacent to the multi-spout cartoon. D) Lick raster of a representative mouse depicting the licks for water, medium concentration, and high concentration during the 3 s access period. E) Mean binned lick rate for all mice for each concentration. (F-G) Cumulative distribution of the number of licks in trials with a ylick (F) and the time of the last lick within each licking bout (G). H) The mean number of licks per trial for each concentration. I) The mean number of licks for each concentration per trial binned by 10 trials over the course of the session. (J-P) same as (C-I), but for data from multi-spout consumption of a gradient of concentrations of quinine. (Q-W) same as (C-I), but for data from multi- spout consumption of a gradient of concentrations of NaCl. (*Main effects listed on plots are results of One-Way RM ANOVA; asterisks depict HSD comparisons between concentrations indicated by horizontal line; Faded lines depict individual mice; see stats table for details)*.

Prior to behavioral training, mice were water-restricted to 80-90% baseline bodyweight (Guo et al., 2014). However, during behavioral sessions, multiple mice were able to consume enough fluid to maintain weight above 90% baseline body weight. Separate groups of mice were used for sucrose, quinine, and sodium chloride solution sets to control for training history. All groups of mice were initially conditioned on free-access licking in 1 - 2 sessions and then conditioned with the multi-spout procedure for 3 - 7 sessions prior to 5 sessions of counterbalanced spout pairing (summarized in **Figure 4**). The licks measured using this approach approximate consumption, as total number of licks during each session is strongly correlated with weight in fluid consumed during the session (**Figure 4- figure supplement 1A**). Using this approach, we successfully elicited a range of consumption responses for each solution set.

Mice displayed gradations in licking for different concentrations of sucrose, quinine, and sodium chloride (**Figure 4C-W**). For each solution set, licking bouts during the access period (representative session depicted in **Figure 4D, K, R**, mean binned lick rate across all trials depicted in **Figure 4E, L, S**) displayed inter-lick intervals similar to freely moving consumption (**Figure 4- figure supplement 1C**). Mice licking for gradations of sucrose (**Figure 4C-I**) showed a modest range of licking behavior where trials with higher concentrations of sucrose elicited a greater number of licks (One-Way RM ANOVA: Concentration***; **Figure 4F, H**) and longer time spent licking during the trial (**Figure 4G**). Mice licking for gradations of quinine (**Figure 4J-P**, **Figure 4-figure supplement 2**) showed a modest range of licking behavior where trials with higher concentrations of quinine elicited a lower number of licks (Loney and Meyer, 2018) (One-Way RM ANOVA: Concentration***; **Figure 4M, O**) and shorter time spent licking during the trial (**Figure 4N**). Mice licking for gradations of NaCl (**Figure 4Q-W**, **Figure 4- Figure Supplement 3**) showed a large range of licking behavior where trials with higher concentrations of NaCl elicited a lower number of licks (One-Way RM ANOVA: Concentration***; **Figure 4T, V**) and shorter time spent licking during the trial (**Figure 4U**). Each solution set produced unique time-courses of licking behavior over the course of the session (**Figure 4H**, **Figure 4- figure supplement 1D**). Mice in the sucrose set started with high licking rates and showed a gradual satiation that resulted in decreased licking across all concentrations (**Figure 4I**, **Figure 4- figure supplement 1D**); mice in the quinine set started with high licking rates but rapidly dropped by around trial 40 across all concentrations (**Figure 4P**, **Figure 4- figure supplement 1D**); and mice in the NaCl set showed only a minor reduction in licking across all concentrations throughout the session (**Figure 4W**, **Figure 4- figure supplement 1D**). Comparing the total number of licks per session across the three sets of solutions revealed that mice displayed the highest number of licks during the sucrose set, then NaCl, then quinine (**Figure 4- figure supplement 1E**). Comparing task engagement using the proportion of trials with licking across the three sets of solutions, mice in the quinine set showed substantially lower proportion of trials with licks compared to mice in the sets for sucrose or NaCl (**Figure 4- figure supplement 1F**). Mice displayed little to no relationship between the number of licks in the session and weight of the mouse or amount of fluid consumed/provided on the previous session (**Figure 4- figure supplement 1G-J**). We also found that older mice displayed higher lick rates for sucrose (**Figure 4- figure supplement 1K-L**). Finally, we found no sex differences in task performance, except a lower proportion of trials with licking in female mice (**Figure 4- figure supplement 4**). Altogether, these data indicate that OHRBETS successfully elicits a range of consumption behavior for differential concentrations of appetitive and aversive solutions.

Given that mice showed a smaller range of licking for gradations in quinine compared to NaCl, we further investigated licking behavior with additional sets of 1:4 serial dilutions of quinine with higher concentrations (starting concentration: Low = 1 mM (**Figure 4** **J-P**), Med = 5 mM, High = 10 mM) (**Figure 4- figure supplement 2**). Each quinine set produced a modest range of licking behavior with less licking for higher concentrations of quinine (**Figure 4- figure supplement 2A-D**). Mice displayed a lower total licking in the High set compared to the Med and Low sets (**Figure 4- figure supplement 2E**), and mice in all sets showed similar task engagement as indicated by proportion of trials with licking (**Figure 4- figure supplement 2F**). Mice in all sets abruptly stopped licking part-way through the session (**Figure 4- figure supplement 2C**). Overall, each quinine set was capable of producing a range of licking behavior but failed to support licking throughout the entirety of the behavioral session.

### Homeostatic demand shifts within-session consumption of gradients of sucrose and NaCl

To determine if OHRBETS multi-spout assay could detect shifts in consumption behavior following behavioral challenges, we measured consumption of a gradient of sucrose concentrations across homeostatic demand states. We trained mice in the multi-spout brief-access task for 5 sessions under water-restriction, then 5 sessions under food-restriction, and ending with 5 sessions under no restriction (*ad-libitum*) (**Figure 5A**). We observed strong effects of restriction state on consumption behavior across sucrose concentrations (**Figure 5B-D**, **Figure 5- figure supplement 1A-E**). Most notably, mice showed a substantially larger range of licking behavior under food-restriction compared to water-restriction and *ad-libitum* (Two-Way RM ANOVA: Concentration x State***; **Figure 5B**). Mice showed vastly different levels of total number of licks with the greatest number of licks for all concentrations under water-restriction, then food-restriction, then *ad-libitum* (Two-Way RM ANOVA: State***; **Figure 5B, C**). Mice also displayed differences in licking rate throughout the session (Two-Way RM ANOVA: State***, Trial Bin x State***; **Figure 5C, D**). The minor scaling in licking across sucrose concentrations under water-restriction compared to food-restriction could indicate that the water component of the solutions is strongly appetitive under water-restriction. Using OHRBETS, we measured changes in the relative consumption of concentrations of sucrose across homeostatic demand states that closely parallels the effect of homeostatic demand on sucrose consumption described in freely moving rodents (Glendinning et al., 2002; Smith et al., 1992; Spector et al., 1998).

**Figure 5:**
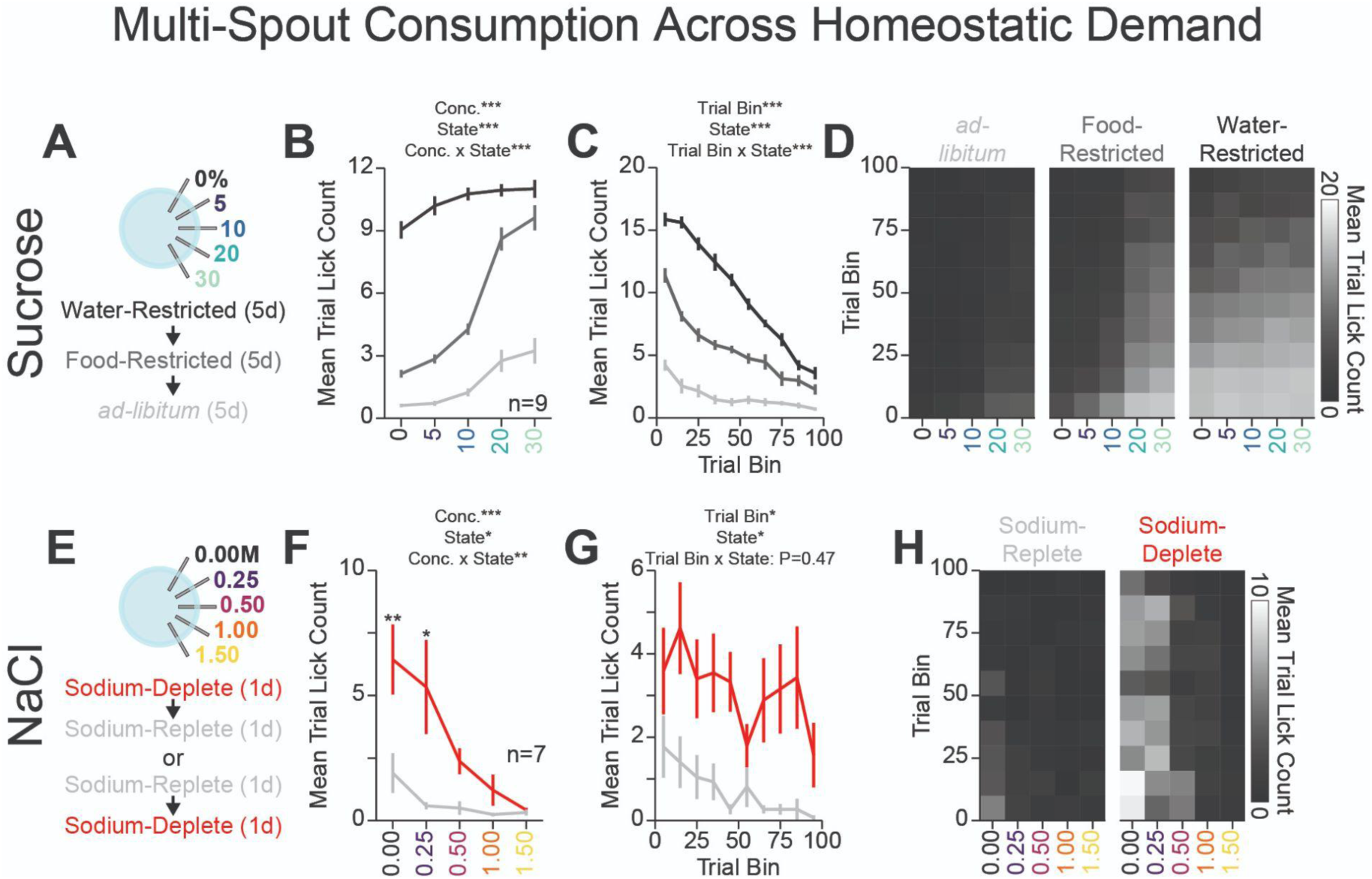
Homeostatic Demand Shifts Within Session Consumption of Gradients of Sucrose and NaCl: A) Procedure: Mice ran sequentially through water-restriction, food-restriction, and *ad-libitum* states During each state, mice received 5 sessions of multi-spout counterbalanced to have each concentration of sucrose on each spout once. B) The mean number of licks per trial for each concentration of sucrose in the *ad-libitum* (light gray), food-restricted (dark gray), and water-restricted (black) states (*HSD: every mean is significantly different from every other, except 30% sucrose consumption under food and water- restriction*). C) Mean trial lick count across all concentrations of sucrose in bins of 10 trials across the session for each homeostatic state. D) The mean number of licks for each concentration of sucrose per trial binned by 10 trials over the course of the session for each homeostatic state. E) Procedure: In sodium replete or sodium deplete states in counterbalanced order, mice received 1 session of multi-spout with a gradient of concentrations of NaCl. The pairing of solution concentrations and spouts remained consistent. F) The mean number of licks per trial for each concentration of NaCl in the sodium replete (gray) and deplete (red) states. G) Mean trial lick count across all concentrations of NaCl in bins of 10 trials across the session for each homeostatic state. H) The mean number of licks for each concentration of NaCl per trial binned by 10 trials over the course of the session for each homeostatic state. (*Main effects listed on plots are results of Two-Way RM ANOVA; asterisks indicate differences between homeostatic demand state at a corresponding concentration; see stats table for details)*.

To determine if our head-fixed multi-spout assay could detect shifts in consumption of NaCl, we measured consumption of a gradient of NaCl concentrations across sodium demand states. We first trained mice under water-restriction (**Figure 4**) before allowing mice to return to *ad-libitum* water. Next, we manipulated sodium appetite using furosemide injections followed by access to sodium depleted chow (sodium-deplete) or standard chow (sodium-replete) and then measured consumption of a gradient of NaCl concentrations in our multi-spout assay over 2 sessions (counterbalanced order of sodium appetite state) (**Figure 5E**). Mice displayed greater licking under the sodium-deplete state compared to the sodium-replete state (Two-Way RM ANOVA: State***, Concentration x State**; **Figure 5F-H**, **Figure 5- figure supplement 1F-J**). Specifically, mice when sodium-deplete showed higher levels of licking for both water and 0.25M NaCl. Mice displayed more licking throughout the session when sodium-deplete, indicating a heightened demand (Two-Way RM ANOVA: State*; **Figure 5G-H**). The increased licking for water when sodium-deplete can potentially be attributed to higher levels of thirst, as previously described (Jalowiec, 1974). Together, these results indicate that mice show a range of consummatory behaviors that are sensitive to homeostatic demand and that OHRBETS offers a platform for assessing shifts in consummatory drive in a reliable fashion in head-fixed mice.

### Light/dark cycle shifts within session consumption of gradients of sucrose

To characterize behavior across the circadian light/dark cycle, we measured consumption of a gradient of sucrose concentrations under food-restriction during the dark cycle or light cycle in separate groups of mice (**Figure 6A**). During 2 sessions of free-access consumption, mice tested in the dark cycle consumed significantly more 10% sucrose compared to mice tested in the light cycle (Two-Way RM ANOVA: Cycle***; **Figure 6B**) (Bainier et al., 2017; Smith, 2000; Tõnissaar et al., 2006). Across 8 sessions of the multi-spout assay, mice tested in the dark cycle licked more compared to mice tested in the light cycle (Cycle**; **Figure 6C** **left**); however, over sessions 4 - 8 there was no effect of light cycle on licking (*t*-test P=0.099; **Figure 6C** **right**). Despite similar overall licking in the multi-spout assay, we found that experiments conducted during the light and dark cycle resulted in distinct licking across sucrose concentrations (Concentration x Cycle***, **Figure 6E**) (Bainier et al., 2017; Tõnissaar et al., 2006). Furthermore, compared to mice tested in the light cycle, mice tested during the dark cycle showed higher levels of consumption early in the session (Time x Cycle***; **Figure 6F, G**). Together, these results indicate that the light/dark cycle affects sucrose consumption and testing mice in the light cycle leads to pronounced reductions in consumption in early training sessions.

**Figure 6:**
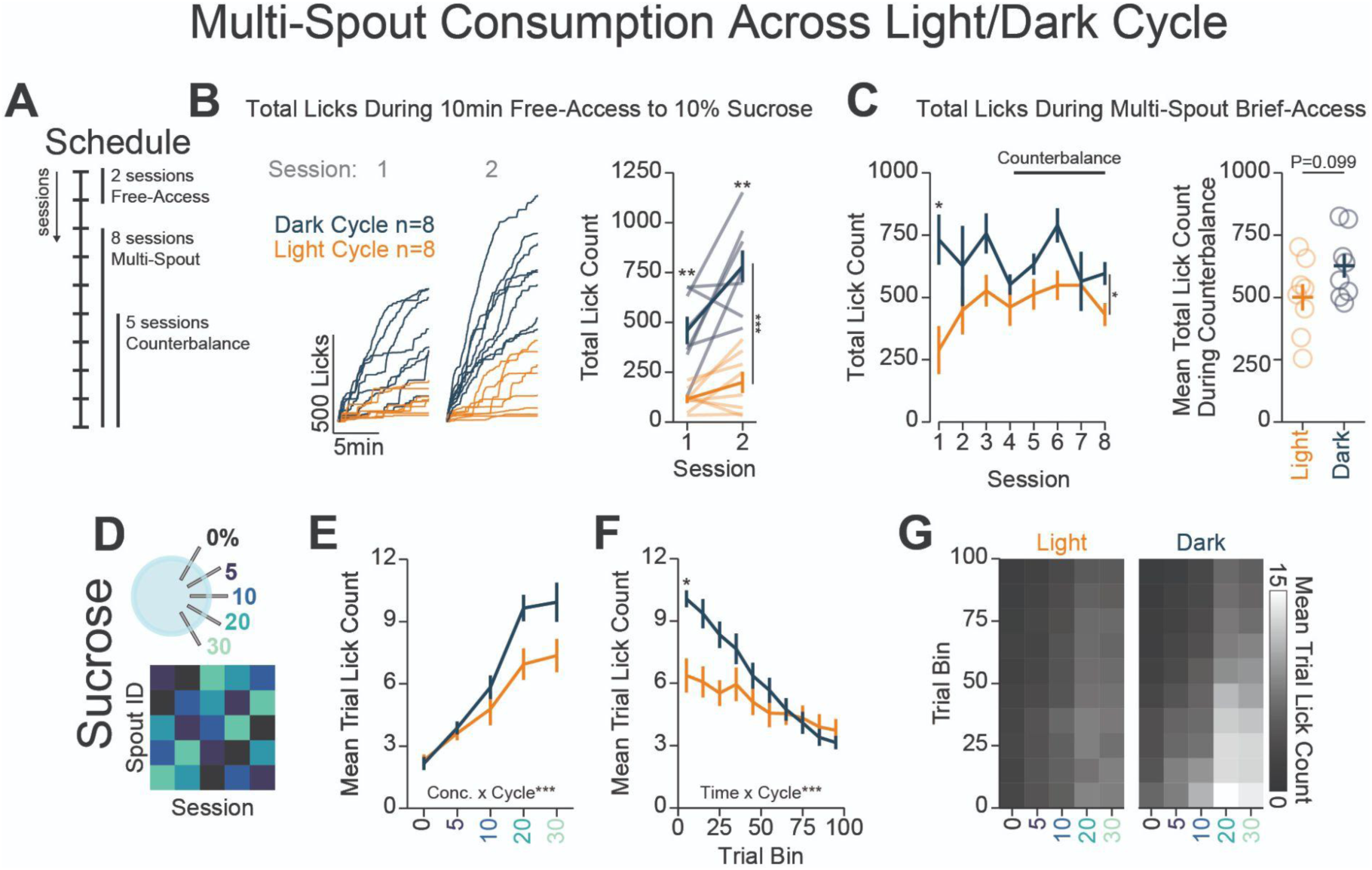
Light/Dark Cycle Shifts Within-Session Consumption of Gradients of Sucrose: A) Schedule for behavioral sessions. B) Licking behavior during two sessions of free-access licking for 10% sucrose displayed as cumulative licking (left) and total lick count during the session (right). Mice ran in the dark- cycle licked more than mice ran in the light-cycle (Two-Way RM ANOVA: Cycle***). C) Total licking behavior during 8 sessions of multi-spout brief-access to a gradient of sucrose concentration (left) and mean over 5 counterbalance sessions (right). Mice ran in the dark-cycle licked more than mice in the light-cycle over all 8 sessions (Two-Way RM ANOVA: Cycle*), but not over the 5 counterbalanced sessions (*t*-test: P=0.099). D) Procedure: mice were trained in 5 sessions of 5x sucrose multi-spout counterbalanced to have each solution of each spout once. E) The mean number of licks per trial for each concentration of sucrose for mice ran in the dark-cycle (blue) and mice ran in the light-cycle (orange) (Two-Way RM ANOVA: Concentration x Cycle***). F) Mean trial lick count across all concentrations of sucrose in bins of 10 trial across the session (Two-Way RM ANOVA: Time x Cycle***). G) The mean number of licks for each concentration of sucrose per trial binned by 10 trials over the course of the session. (*Asterisks above means indicate differences between mice tested in each cycle during the same session*).

### Comparing the reproducibility of the multi-spout brief-access task across independent laboratories

To determine if our system produces quantitatively similar consumption across labs, we compared behavior of food-restricted mice tested in the dark cycle trained on the multi-spout brief-access to a gradient of sucrose concentrations obtained with our head-fixed system across independent labs and geographic locations (**Figure 6- figure supplement 1**; data collected in the Stuber lab is shown in **Figure 5**, and data collected in the Roitman lab is shown in **Figure 6**). We observed qualitative differences in the binned licking rate over the 3 s access period (**Figure 6- figure supplement 1B**), with higher licking rate in mice tested in the Roitman lab near the onset of the access-period. We also found that mice tested in the Roitman lab exhibited a small, but significant, reduction in inter-lick intervals compared to the Stuber lab (**Figure 6- figure supplement 1**). However, despite these nominal differences, there were no statistical differences in the mean licking for each concentration of sucrose across labs (**Figure 6- figure supplement 1D**). These data indicate that our system produces similar consumption behavior when run in different labs, geographic locations, and experimenters.

### OHRBETS combined with fiber photometry to assess ventral striatal dopamine dynamics to multiple concentrations of rewarding and aversive solutions

To demonstrate the utility of the multi-spout assay run on OHRBETS, we performed simultaneous dual fiber-photometry in the mesolimbic dopamine system during the multi-spout assay. The activity of ventral tegmental area dopamine neurons and the release of dopamine in the nucleus accumbens are well known to scale with relative reward value such that the most rewarding stimuli produces increases in dopamine release and the least rewarding stimuli produces modest decreases in dopamine release (Eshel et al., 2015; Hajnal et al., 2004; Tobler et al., 2005). We used multi-spout brief-access to a gradient of an appetitive solution (sucrose) and an aversive solution (NaCl) to elicit a range of consummatory responses (**Figure 4, 5**) while simultaneously recording dopamine dynamics in the medial nucleus accumbens shell (NAcShM) and lateral nucleus accumbens shell (NAcShL) (**Figure 7A**, placements shown in **Figure 7- Figure Supplement 6**). To record dopamine dynamics in the NAc, we expressed the dopamine sensor GRAB-DA (GRAB-DA1h (Sun et al., 2018) or GRAB-DA2m (Sun et al., 2020)) in the NAcShM and NAcShL (counterbalanced hemispheres across mice) of wild-type mice and implanted bilateral optic fibers with a head-ring to facilitate head-fixation (**Figure 7A**, *Methods*). Mice were tested with multi-spout access to a gradient of sucrose concentrations under water-restriction and food- restriction, in counterbalanced order, and then a gradient of NaCl concentrations under water-restriction (**Figure 7B**). Across each stage of the task, mice exhibited scaling in licking behavior that replicated data shown in **Figure 4** and **Figure 5** (**Figure 7-figure supplement 1**). During the multi-spout assay, we observed dynamics in dopamine signals in both the NAcShM and NAcShL during the consumption access period (**Figure 7C-O**). During consumption of sucrose under food-restriction, where we observe a large range in licking across concentrations of sucrose (**Figure 7- figure supplement 1, left**), we measured strong scaling of GRAB-DA fluorescence in the NAcShL and moderate scaling in the NAcShM (representative mouse **Figure 7D**; mean fluorescence **Figure 7E**; mean fluorescence during access **Figure 7F**, Two-Way RM ANOVA: Solution x Region***; CDF shown in **Figure 7-figure supplement 2A**). Specifically, we observed significantly higher responses in the NAcShL compared to the NAcShM at higher concentrations of sucrose (10, 20, and 30% HSD*). On a trial-by-trial basis, we observed a correlation between the amount of licking on a trial and GRAB-DA fluorescence (**Figure 7G**; CDF shown in **Figure 7-figure supplement 2B**). During consumption of sucrose under water-restriction, where we observe high levels of licking but minimal range across concentrations of sucrose (**Figure 7-figure supplement 1, mid**), we measured moderate scaling of GRAB-DA fluorescence in the NAcShL and little scaling in the NAcShM (representative mouse **Figure 7H**; mean fluorescence **Figure 7I**; mean fluorescence during access **Figure 7J**, Two-Way RM ANOVA: Solution x Region***; CDF shown in **Figure 7-figure supplement 2C**). Specifically, mice displayed significantly higher GRAB-DA responses in the NAcShL compared to the NAcShM at higher concentrations of sucrose (**Figure 7F**). Like dynamics during food-restriction, GRAB-DA fluorescence was positively correlated with licking within the trial (**Figure 7K**, CDF shown in **Figure 7-figure supplement 2D**). During consumption of the aversive tastant (NaCl) under water-restriction, where we observed a large range of licking across concentrations of NaCl (**Figure 7-figure supplement 1, right**), we measured strong scaling of GRAB-DA fluorescence in both the NAcShL and NAcShM (representative mouse **Figure 7L**, mean fluorescence **Figure 7M**; mean fluorescence during access **Figure 7N**, Two-Way RM ANOVA: Solution x Region***, CDF shown in **Figure 7-figure supplement 2E**). Despite the interaction between solution and region of the NAc, there was a significantly higher GRAB-DA fluorescence in the NAcShL only during 0.25M NaCl. Like other stages of the task, we observed a clear correlation between GRAB-DA fluorescence and licking during the trial (**Figure 7O**, CDF shown in **Figure 7- figure supplement 2F**). Taking advantage of the head- fixed preparation, we were able to record the activity of the NAcShL and NAcShM simultaneously and found a strong correlation in GRAB-DA fluorescence in the two regions across each stage of the task (**Figure 7-figure supplement 3**).

**Figure 7:**
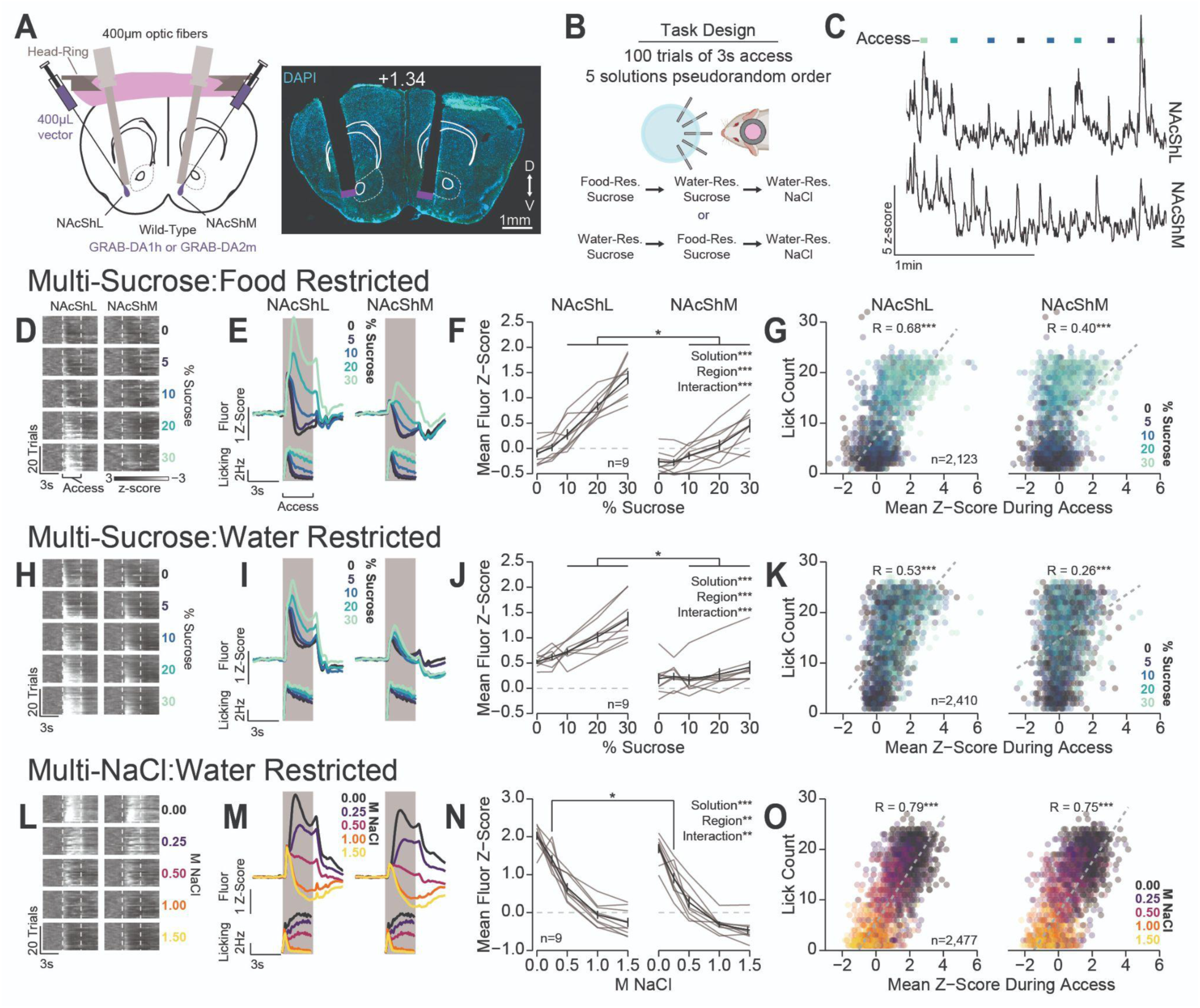
Differential Dopamine Dynamics During Multi-Spout Consumption Behavior: A) Approach for simultaneously recording dopamine dynamics in the NAcShL and NAcShM (left), and representative placements of optic fibers overlaying the NAcSh (White numerical value indicates AP position relative to bregma). B) Task design and schedule of experiment. C) Representative trace of simultaneous GRAB- DA fluorescence in the NAcShM and NAcShL during multi-spout access to sucrose under food-restriction (lines on top indicate access periods, color indicates sucrose concentration). (D-G) Dopamine dynamics during multi-sucrose under food-restriction: D) Representative heat map of GRAB-DA fluorescence over time during each trial sorted by sucrose concentration (trials averaged over 3 sessions of recording). E) Perievent time histograms of GRAB-DA fluorescence (top) and licks (bottom). F) Mean fluorescence *z*- score during access period indicating strong scaling in the NAcShL (left) and weak scaling in the NAcShM (right). G) On individual trials, the mean *z*-score during access correlates with licking in both the NAcShL (left), and NAcShM (right) (color depicts the solution concentration). (H-K) Same as D-G for dopamine dynamics during multi-sucrose under water-restriction. (L-O) Same as D-G for dopamine dynamics during multi-NaCl under water-restriction. N) Strong scaling in NAcShL (left) and weak scaling in the NAcShM (right). (*Asterisks indicate differences between brain regions at the same solution concentration; Two- Way RM ANOVA effects indicated in F, J, N; correlations indicated in G, K, O; see stats table for details*).

Interestingly, the NAcShM and NAcShL show a differential range of GRAB-DA fluorescence across each stage of the task. The NAcShM shows a disproportionately higher range of GRAB-DA fluorescence during multi-spout consumption of NaCl compared to the other stages of the task (**Figure 7-figure supplement 4B**, Two-Way RM ANOVA: Stage x Brain Region***). The higher range of dopamine release in the NAcShM during consumption of a range of aversive solutions compared to appetitive solutions could indicate a specialized role for the NAcShM in mediating behavioral responses to aversive stimuli as previously described for footshock conditioning (Jong et al., 2018). OHRBETS allowed us to isolate consumption behavior in response to a range of rewarding and aversive solutions while performing dual site fiber photometry and revealed robust dopamine responses that scales with solution value and consumption. Furthermore, these data indicate that OHRBETS is highly compatible with neural recording and manipulation techniques that would be challenging with freely moving behavioral designs.

## Discussion

OHRBETS is a customizable, inexpensive system for head-fixed behavior in mice that enables a variety of behavioral experiments, including operant conditioning, real-time place testing, and multi- solution brief-access consumption, accurately replicating behaviors in freely moving. These data demonstrate that a diverse set of operant and consummatory behaviors are compatible with head-fixed procedures run with a single hardware setup and will serve as a resource for future investigations into these behaviors using neuroscience approaches that rely on head-fixation.

Behavior measures within our head-fixed adaptations of freely moving operant assays reproduce many important phenotypes originally characterized in freely moving behavior. Mice rapidly learn operant responding for sucrose and then flexibly express responding as a function of reward-cost and reward- size (Kliner et al., 1988; Reilly, 1999; Sclafani and Ackroff, 2003; Winger and Woods, 1985). Using optogenetic stimulation, which offers tighter control over the precise magnitude and timing of appetitive and aversive states, mice exhibited quantitatively similar positive ICSS, RTPP, and RTPA behavior with our head-fixed and freely moving approaches. The ability to conduct ICSS and RTPT with a single setup is particularly useful in measures of valence-related neural circuits, but these results also imply that the head-fixed RTPT procedure could be used to test the appetitive and aversive quality of other stimuli that are challenging to test in freely moving conditions including discrete somatosensory stimuli. Taken together, our results establish that our behavioral system produces robust, reproducible operant behavior consistent with the commonly employed freely moving counterparts.

In addition to the operant conditioning experiments, our system can facilitate multi-solution brief- access experiments for studying consummatory behavior. In our task, mice show consumption of a gradient of sucrose, quinine, and NaCl concentrations that closely matches behavior with the freely moving version of the task (Corbit and Luschei, 1969; Coss et al., 2022; Garcia et al., 2021; Glendinning et al., 2002; John et al., 1994; Loney and Meyer, 2018; Smith et al., 1992; Villavicencio et al., 2018). Licking increased monotonically with increased concentrations of sucrose across all homeostatic states (Garcia et al., 2021; Glendinning et al., 2002; Smith et al., 1992; Spector et al., 1998). However, homeostatic demand states produced pronounced differences in the range of consumption behavior across sucrose concentration, as food-restriction produced a substantially larger range of licking behavior compared to water-restriction. One unexpected finding was that mice showed vastly different behavior when licking for the aversive tastants quinine and hypertonic NaCl. When licking for quinine, mice abruptly ceased consumption for all concentrations mid-way through the session. On the other hand, when licking for NaCl, mice continue to consume large amounts of low concentrations of NaCl throughout the entire session. These results may be explained by an additive effect of quinine that builds in aversion over trials and results in a lingering bitter taste (Leach and Noble, 1986) that attenuates motivation to initiate consumption. During the NaCl sessions, NaCl may stimulate thirst (Kraly et al., 1995; O’Kelly, 1954; Stricker et al., 2002) resulting in enhanced motivation to consume water. Thus, the multi-spout brief access task with gradients of NaCl can be a uniquely advantageous approach for eliciting a high number of strongly aversive events while continuing to engage behavior. Changes in task design could improve performance during the quinine task, such as including water rinse trials between each quinine trial (Loney and Meyer, 2018). In addition to using a gradient of solution concentrations, any number of combinations of tastants could be used to study a whole host of behavioral phenomena including innate and conditioned consumption behaviors.

Using our multi-spout brief-access task in conjunction with GRAB-DA fiber-photometry, we observed dopamine dynamics that positively correlated with relative solution value and consumption. Previous studies have revealed that dopamine release in the ventral striatum (Hajnal et al., 2004) and dopamine neuron activity scales with reward magnitude (Eshel et al., 2015; Tobler et al., 2005). We found that dopamine release in these subregions’ scales with the relative value of the solution being consumed and the amount of concurrent consumption and is strongly influenced by solutions present in a session and the mouse’s homeostatic demand state. Most interestingly, we found that the dopamine release in the NAcShM has a much larger amplitude during multi-spout consumption of a gradient of NaCl concentrations than during consumption of a gradient of sucrose concentrations. This result implies that dopamine release in the NAcShM tracks value, and the range of values during the multi-spout consumption of gradients of NaCl is greater than the range of values during multi-spout consumption of gradients of sucrose. Alternatively, this result could indicate a specific role of dopamine release in the NAcShM that corresponds to shaping behavior or learning in the face of aversive events (Jong et al., 2018). By conducting these experiments using OHRBETS, we removed approach behaviors that occur prior to consumption and isolated neuronal responses specifically during consumption (Chen et al., 2022) without interference of activity ramps observed in freely moving behavioral designs (Howe et al., 2013). Future experiments are necessary to reveal the specific contribution of licking, taste, and value to wide- spread dopaminergic signals and how these signals causally influence ongoing consumption or learning.

Eliminating locomotion improves compatibility with many standard neuroscience approaches including optogenetics, fiber-photometry, electrophysiology, and calcium imaging. To prevent twisting of tethers, each of these approaches require a commutator in freely moving conditions, but with head- fixation the need for a commutator is eliminated. This facilitates multiplexed experiments with simultaneous use of multiple approaches that each rely on independent tethers without the risk of weighing down the animal, tangling, or twisting to the point of affecting task performance. For fiber- photometry, fixing the animal dramatically reduces motion artifacts, thereby reducing the need for an isosbestic to correct for motion (**Figure 7- Figure Supplement 5**). This opens the ability to conduct experiments with fluorescence biosensors without known isosbestic points or without true isosbestic points. Recent advances in optical imaging have opened up new approaches in freely moving animals, but the cutting edge of optical technologies will typically start with tabletop microscopes. Using head- fixed models permits users to embrace cutting edge imaging technologies without waiting for further advances to bring the technology into freely moving animals. The use of OHRBETS allows for enhanced compatibility with a variety of neuroscience technologies and will enable novel, multiplexed experiments that would be difficult or impossible to conduct in freely moving animals.

While head-fixed experiments offer many advantages, they come with important caveats, limitations, and experimental design considerations. Head-fixation can be acutely stressful to mice and causes increased levels of circulating stress markers (Juczewski et al., 2020), which could impair learning and interact with other manipulations. The advantage of limiting the range of behaviors a subject can display comes at the cost of reduced naturalistic character, which can impair behavior and related neuronal activity (Aghajan et al., 2015; Aronov and Tank, 2014). Furthermore, isolation of components of behavior provides powerful insight into the neuronal mechanisms that underlie the particular component of behavior but may impair insight into how the related neuronal circuits function during more complex behaviors and contexts. Even in the presence of these caveats, extensive research conducted in head-fixed non-human primates has made vast progress in a multitude of areas of neuroscience (Mirenowicz & Schultz, 1996; Parker & Newsome, 1998; Schultz et al., 1997) including appetitive and consummatory behaviors (Bromberg-Martin et al., 2010; Haber & Knutson, 2010). The greatest insights into the neuronal mechanisms of behavior will come from a mixture of both naturalistic behaviors and highly controlled behaviors facilitated by head-fixed behaviors made possible with OHRBETS.

The OHRBETS ecosystem presented here was designed to be scalable, flexible, and compatible with external hardware. By using low-cost, open-source, and 3D printed components and publishing extensive instructions for assembly, our system is affordable and scalable across labs of all sizes and budgets. Despite the use of low-cost and 3D printed components, our system is remarkably consistent and reliable across hundreds of behavioral sessions. Our hardware and software are modular, as all hardware components can be easily swapped, and all behavioral programs are written to produce data with a uniform format. Using different combinations of components will facilitate conducting a wide variety of behavioral experiments including all the experiments presented in this manuscript and many more. By using an Arduino Mega case as a microprocessor mounted within a 3D printed enclosure, one can integrate many different forms of connectivity to interface with external hardware. In the online models, we have options for communication via BNC, Cat6, and DB25 that can be easily combined to suit the user’s needs. Altogether, OHRBETS is a complete platform for diverse behavioral experiments in head- fixed animals that can be easily adapted by the broader scientific community to conduct an even wider range of procedures that are compatible with monitoring and manipulating neural dynamics *in vivo*.

## Materials and Methods

### Instructions for Assembling the OHRBETS

Detailed part list, 3D models, electronic wiring diagrams, behavioral programs, and instructions for assembling are available publicly on our GitHub repository (https://github.com/agordonfennell/OHRBETS).

### Hardware

3D printed components designed and available via the web-based cad software TinkerCAD (Autodesk) and printed using a filament printer (Ultimaker S3) using PLA or resin printer (Form3) using Clear Resin. We also tested components built by the online printing service CraftCloud and found that they work similarly to ones printed in the lab. The micropositioner design was based on one created by Backyard Brains (Backyard Brains, 2013) and the retractable spout design was based on one created by an independent designer (Buehler, 2016b).

All behavioral hardware was controlled using an Arduino Mega 2560 REV3 (Arduino). The timing of events was recorded via serial communication from the Arduino to the computer (PC, running Windows10) by USB. Lick spouts were made by smoothing 23 gauge blunt fill needles using a Dremel with a sanding disk. Liquid delivery was controlled by solenoids (Parker) gated by the Arduino, using a 24V transistor. The retractable spout, radial spout, and wheel brake utilized micro servos (Tower Pro SG92R). Licks on each spout were detected individually using a capacitive touch sensor (Adafruit MPR121) attached to each metal spout. Importantly, the baseline capacitance of each sensor was kept to a minimum and touch thresholds were reduced from standard values (see GitHub for detailed instructions). Micropositioners were assembled from 3D printed components, Super Glue (Loctite Super Glue ULTRA Liquid Control), screws, and nuts.

### Hardware Validation

We measured the consistency of the retractable spout extension latency and terminal positions using video recording. We recorded 1,000 extension/retractions in 5 separate retractable spouts using a high-speed video camera (Basler, acA800-510um, 200 fps). We then estimated the position of the spout using DeepLabCut (Mathis et al., 2018) and analyzed the position of the spout relative to the mean terminal position of the spout over time. We measured the consistency of the wheel brake latency using experimenter and mouse rotation. We recorded 1,770 wheel rotations produced by an experimenter and measured the effect of braking using 4 separate head-fixed systems. We computed the binned rotational velocity by taking the mean instantaneous velocity within 25ms time bins (**Figure 1E**, **Figure 1-Figure Supplement 1 H**). We also assessed the rotation following brake engagement with all brake events during all operant data included in **Figure 1**.

### Software

All behavioral programs were written in the Arduino language and executed on the Arduino Mega during the behavioral session. The timing of hardware and behavioral events were sent from the Arduino and recorded on a PC computer (Windows 10) via serial communication or through a fiber-photometry console via TTL communication. Fiber-photometry data was collected using Synapse (Tucker Davis Technologies). Data processing, statistical analysis, and data visualization was performed using custom scripts in Python (version 3.7) and R (version 4.0.4). All behavioral programs and pre-processing scripts used to produce the data in this manuscript are freely available through our GitHub (https://github.com/agordonfennell/open_stage).

### Animals

All behavioral procedures were pre-approved by University of Washington or University of Illinois at Chicago Animal Care and Use Committees. A mixture of wild-type and transgenic mice on a C57BL/6J background were used for experiments throughout the paper. All mice were bred in the lab from mouse lines obtained from Jackson Laboratory aside from 16 wild-type mice obtained directly from Jackson Laboratory. No differences were observed across transgenic lines so all data was pooled. Mice used in fiber-photometry and optogenetic experiments were singly housed to prevent damage to the optical fibers while all other mice were group housed. Mice were kept on a reverse 12h light/dark cycle and behavioral experiments were conducted within the dark-cycle unless otherwise noted.

### Surgeries

Mice (>P55) were anesthetized using isoflurane (5% induction, 1.5-2% maintenance), shaved using electric clippers, injected with analgesic (carprofen, 10 mg/kg, s.c.), and then mounted in a stereotaxic frame (Kopf) with heat support. Skin overlying the skull was injected with a local anesthetic (lidocaine, 2%, s.c.) and then sterilized using ethanol and betadine. Next, an incision was made using a scalpel, and the skull was cleared of tissue and scored using the sharp point of a scalpel. The skull was leveled, 2 burr holes were drilled in the lateral portion of the occipital bone, and 2 micro screws were turned into the bone. We then coated the bottom of a stainless-steel head-ring (custom machined, see GitHub for design) with Super Glue, placed it onto the skull of the mouse, and then encased the head- ring and skull screws with dental cement making sure the underside of the ring remained intact. After the dental cement had time to fully dry, the mouse was removed from the stereotaxic frame and allowed to recover with heat support before being returned to their home cage. Mice were allowed to recover for at least 1 week prior to dietary restriction.

Mice used for optogenetic or fiber-photometry experiments underwent the same procedure as above with the addition of a viral injection and fiber implantation. Following implantation of skull screws, we drilled a burr hole overlaying the brain region target. We then lowered a glass injection pipette into the target brain region and injected the virus at a rate of 1nL/s using a Nanoject III (Drummond), waited 5 min for diffusion, and then slowly retracted the pipette. For optogenetic experiments, we injected 300nL of AAV5-EF1a-DIO-hChR2(H134R)-eYFP (titer: 3.2e12) or AAV5-Ef1a-DIO-mCherry (titer: 3.3e12), and for fiber-photometry experiments, we injected 400nL of AAV9-hSyn-GRAB-DA1h (titer: 2.7e13) or AAV9- hSyn-GRAB-DA2m (titer: 2.4e13). The following stereotaxic coordinates (relative to bregma) were used for injection targets: LHA (0° angle; AP: -1.3 mm; ML: +/- 1.1 mm; DV: -5.2 mm), NAc medal shell (10° angle; AP: 1.7 mm; ML: +/- 1.5 mm; DV: -4.8 mm), and NAc lateral shell (10° angle; AP: 1.7 mm; ML: +/- 2.5 mm; DV: -4.6 mm). Next, we lowered a 200 µm optic fiber for optogenetic experiments (1.25mm ferrule, 6mm fiber length, RWD) or a 400 µm optic fiber for fiber-photometry experiments (2.5 mm ferrule, 6 mm fiber length, MFC_400/470-0.37_6mm_MF2.5_FLT, Doric) 0.2 mm dorsal to the injection site and then encased the fiber extending from the brain, metal ferrule, and head-ring with Super Glue and dental cement. Mice were allowed to recover for at least 2 weeks prior to behavior or dietary restriction.

### Behavior

#### Habituation to Head-Fixation and Free-Access Lick Training

Prior to head-fixed behavior, mice were habituated to the experimenter and head-fixation stage over 4 sessions. In the first session, mice were brought into the behavioral room and allowed to explore the head-fixed apparatus to become acquainted with the sights, smells, and sounds of the behavioral box. On the second session, mice were brought into the behavior room and scruffed twice. In the third session, mice were brought into the behavior room, scruffed twice, and then gently had their rear end and hind paws placed in a 50 mL conical twice. After each habituation session, the mouse was immediately provided food or water depending on their deprivation status. Mice undergoing head-fixed operant conditioning for sucrose or head-fixed multi-spout consumption were habituated to head fixation and trained to lick for sucrose in a single 10 min session. During this session, mice were given free- access to water (mice used for quinine and NaCl multi-spout experiments Figure 4,5) or 10% sucrose (mice used for all other experiments). Free-access was approximated using closed-loop delivery of a pulse of fluid (∼1.5 µL) each time the mouse licked the spout. During the training session, mice were head-fixed, and the spout was brought forward to gently touch the mouse’s mouth to encourage licking before being moved to be positioned ∼2-3 mm in front of the mouse’s mouth where it remained throughout the session.

### Head-fixed Operant Conditioning for Sucrose

Retractable spout training consisted of 3 daily sessions of 60 trials with 5 s access periods separated by 20-40 s inter-trial intervals. During each access period, an auditory tone (5 kHz) was played and 5 pulses of ∼1.5 µL sucrose were delivered with a 200ms inter pulse interval. We delivered pulses of sucrose to encourage licking in bouts and to minimize the chance that a large droplet of sucrose would fall.

Operant conditioning training consisted of 6 30 min sessions of initial training, 4 sessions of increased fixed-ratio, and then 5-6 sessions of progressive-ratio. Throughout operant conditioning, one direction of rotation was assigned as the active direction and the opposite direction was assigned as inactive (counterbalanced across mice). Rotation in the active direction earned sucrose delivery. Each sucrose delivery consisted of wheel brake engagement, followed by spout extension and 5 pulses of ∼1.5 µL of 10% sucrose with an inter pulse interval of 200ms. During the 3 s access period, the spout remained extended, a 5 kHz auditory tone was presented, and the brake was left engaged. Rotation in the inactive direction led to wheel brake engagement for the same length of time as the total brake time with active rotation, but the spout did not extend, and sucrose was not delivered. During initial training, the fixed- ratio of reward was 1/4 turn in the first session and 1/2 turn during the next 5 sessions. During increased cost sessions, the fixed-ratio was increased to 1 turn. A total of 3 mice that underwent initial training were removed from progressive ratio training, 2 for not learning the task and 1 that lost their headcap during behavior. During progressive ratio, the wheel turn cost of reward was increased either semilogarithmic (0.25, 0.5, 0.81, 1.21, 1.71, 2.3, 3.1, 4.1, etc., approximating (Richardson and Roberts, 1996) or linearly by 0.5 rotation (0.5, 1.0, 1.5, 2.0, 2.5, 3.0, etc.) each time a reward was earned. The session duration was 1h, or 15 min without earning a reinforcer, whichever comes first.

Reversal training was performed in a naive cohort of mice using an identical procedure to initial operant conditioning training, expert in session 6 th direction of the wheel rotation that was reinforced was inverted (right turn reinforced -- left turn reinforced). Following reversal, the mice were trained on the task for an additional 7 sessions of operant conditioning.

### Optogenetic Experiments

*Vgat*-cre and *Vglut2*-cre mice underwent surgery for experiments with optogenetics outlined above. After at least 4 weeks of recovery, mice were trained on head-fixed and freely moving versions of RTPT and ICSS experiments in series. All mice were trained on RTPT prior to ICSS, but the order of head-fixed and freely moving versions were counterbalanced across mice.

Freely moving RTPT consisted of 1 session of habituation and 6 sessions of RTPT with different stimulation frequencies. During habituation, mice were scruffed, attached to an optic fiber, and allowed to explore the RTPT chamber for 10 min. The RTPT chamber was a two-chamber apparatus (50 × 50 × 25 cm black plexiglass) with two identical compartments. Over the next 6 sessions, mice underwent daily 20 min RTPT sessions with stimulation paired with one of the two compartments. The position of each mouse was tracked in real time using Ethovision (Noldus) and when the mouse’s center point was detected in one of the two compartments it triggered continuous laser stimulation (5ms pulses, ∼10mW power, frequencies: 0, 1, 5, 10, 20, 40 Hz). To prevent associations between stimulation and chambers in the RTPT chamber, the stimulation frequency and compartment paired with laser stimulation were counterbalanced across sessions.

Head-fixed RTPT consisted of 3 sessions of habituation and 12 sessions of RTPT with different stimulation frequencies (6 sessions without and 6 sessions with a tone indicating the mouse’s position). During habituation, mice were habituated to head-fixation as outlined above but without sucrose provided. Over the next 12 sessions, mice underwent daily 20 min RTPT sessions with stimulation paired to one half of the wheel. Throughout RTPT, mice were head-fixed, an optic fiber was connected and covered using blackout tape, and the start of the session was indicated when the wheel brake was disengaged. At the start of the session, the starting wheel position was set in the unpaired zone adjacent to the paired zone. The wheel rotation was tracked by recording the rotation of the wheel relative to the starting position (64 positions/1 rotation) and when the mouse’s position was detected in one of the two zones it triggered continuous laser stimulation (5ms pulses, ∼10mW power, frequencies: 0, 1, 5, 10, 20, 40 Hz). During sessions 1-6 of RTPT, there were no extraneous cues indicating which zone the mouse was located in. During sessions 7-12 of RTPT, there were tone cues (5 and 10 kHz) that indicated if the mouse was in the paired or unpaired zones of the wheel. To prevent learned associations between stimulation and zones over multiple sessions, we counterbalanced the following factors across sessions: the stimulation frequency, side of the wheel paired with laser stimulation, and the tone paired with laser stimulation.

Freely moving operant conditioning for optogenetic stimulation with positive reinforcement consisted of 1 session of habituation, 4 sessions of ICSS training, and 6 sessions of ICSS with different stimulation frequencies. During habituation, mice were scruffed, attached to an optic fiber, and allowed to explore the ICSS chamber (MED Associates) for 20 min. The ICSS chamber contained 2 nose pokes with a light cue located inside and a light cue located above each nose poke, as well as an auditory tone generator. Time stamps of hardware and behavioral events were recorded using MED associates. During daily 20 min ICSS sessions, 1 nose poke into the active nose poke triggered 1 s of laser stimulation and concurrent illumination of the active nose poke light cues and 5 kHz auditory tone. Nose pokes during the 1 s stimulation period were recorded but did not result in an additional stimulation. Nose pokes in the inactive hole were recorded but had no programmed consequence. To train mice to respond for laser stimulation, mice were run through 5 sessions of ICSS training with 20 Hz stimulation. To measure the operant response rates across stimulation frequencies, mice were run though an additional 6 sessions of ICSS with different stimulation frequencies (5ms pulses, ∼10mW power, frequencies: 0, 1, 5, 10, 20,

40 Hz).

Head-fixed operant conditioning for optogenetic stimulation with positive reinforcement consisted of 5 sessions of ICSS training and 6 sessions of ICSS with different stimulation frequencies. During daily 20 min ICSS sessions, wheel rotation in the active direction triggered 1 s of laser stimulation and concurrent 5 kHz auditory tone. The wheel brake was disengaged throughout the behavioral session. Rotation during the 1 s stimulation period was recorded but did not count towards additional stimulation. Rotation in the inactive direction was recorded but had no programmed consequence. Mice were trained over 5 sessions of ICSS for 20 Hz stimulation with a fixed-ratio of 1/4 turn on session 1 and fixed-ratio of 1/2 turn on sessions 2-5. To measure the operant response rates across stimulation frequencies, mice were tested over an additional 6 sessions of ICSS with different stimulation frequencies (5ms pulses, ∼10mW power, frequencies: 0, 1, 5, 10, 20, 40 Hz) and a fixed-ratio of 1/2 turn.

Freely moving operant conditioning for optogenetic stimulation with negative reinforcement consisted of 1 session of habituation and 3 sessions of ICSS training. The habituation and behavioral hardware were identical to the freely moving operant conditioning for optogenetic stimulation with positive reinforcement described above. During daily 20 min ICSS sessions, continuous stimulation was turned on at the start of the session and 1 nose poke into the active nose poke paused laser stimulation for 3 s and triggered concurrent illumination of the active nose poke light cues and 5 kHz auditory tone. Nose pokes during the 3 s pause period were recorded but did not result in an additional pause. Nose pokes in the inactive hole were recorded but had no programmed consequence.

Head-fixed operant conditioning for optogenetic stimulation with negative reinforcement consisted of 11 sessions of ICSS training. During daily 20 min ICSS sessions, laser stimulation was turned on at the start of the session and wheel rotation in the active direction paused laser stimulation for 3 s and triggered a concurrent 5 kHz auditory tone. The wheel brake was disengaged throughout the behavioral session. Rotation during the 3 s pause period was recorded but did not count towards additional pause. Rotation in the inactive direction was recorded but had no programmed consequence. Mice were run through 5 sessions of ICSS training for 5 Hz stimulation and then 6 sessions for 10 Hz stimulation with a fixed-ratio of 1/4 turn in session 1 and fixed-ratio of 1/2 in sessions 2-11.

### Head-fixed Multi-Spout Consumption

Mice were habituated to head-fixation as outlined above, and then ran through 1-3 sessions of spout training (see *Habituation to Head-Fixation*). The multi-spout assay consisted of daily sessions with 100 trials of 3 s access to 1 of 5 different solutions (pseudorandom order with 2 presentations of each solution per every 10 trials), each session with an inter-trial interval of 5-10 s sampled from a uniform distribution. Licks were detected using a capacitive touch sensor and triggered solution delivery via solenoid opening. The duration of opening for each solenoid was calibrated before each experiment to deliver approximately 1.5 µL per solenoid opening by weighing the weight of water produced with 100 solenoid openings. To control for a spout effect, the pairing of solutions to spouts was counterbalanced such that each spout was paired with each solution over every 5 sessions. Mice were trained for a minimum of 3 sessions prior to the 5 consecutive sessions that are averaged together and used for analysis. Mice in experiments with quinine or NaCl were initially trained in the multi-spout assay with water on all 5 spouts for 3 sessions prior to introducing quinine or NaCl solutions. Mice in quinine experiment were tested with the low quinine dilution set (1 mM, 1:4 serial dilution) for 8 sessions, high (10 mM, 1:4 serial dilution) for 3 sessions, and med (5 mM, 1:4 serial dilution) for 3 sessions in series.

### Homeostatic Demand Multi-Spout Experiments

For experiments with alterations in the homeostatic demand for sucrose solution, mice were trained on the multi-spout assay under 3 homeostatic states in series (water-restricted, food-restricted, then *ad-libitum*). First, mice under water-restriction were trained in the multi-spout assay for different concentrations of sucrose over 8 sessions. Mice were then removed from water-restriction and maintained on food-restriction for 1 week prior to being run through the multi-spout assay for 5 sessions. Finally, mice were removed from all restrictions and maintained with *ad-libitum* access to food and water for 3 sessions prior to being run through the multi-spout assay for 8 sessions. The final 5 sessions from each homeostatic demand state were used for analysis. Mice that went through the fiber-photometry recording experiment were run through the same procedure except the order of water-restriction and food-restriction was counterbalanced across mice, and they were run for 3 sessions of free-access spout training.

For experiments with alterations in the homeostatic demand for sodium chloride, mice were trained under water-restriction and were then run under two homeostatic states in counterbalanced order (sodium-deplete, sodium-replete). First, mice under water-restriction were trained in the multi-spout assay with different concentrations of sodium chloride over 10 sessions. Mice were removed from water-restriction and given *ad-libitum* access to water for 48h prior to manipulations of sodium demand. To generate sodium demand, we used 2 injections of diuretic furosemide (50mg/kg) over 2 days (Jarvie and Palmiter, 2017). Mice were weighed, injected with furosemide, and then placed into a clean cage with bedding for 2 hours before being weighed again to confirm diuretic effect (∼5% weight loss). Mice were then returned to a clean home cage with *ad-libitum* access water and sodium free chow (Envigo, TD.90228) (sodium-deplete) or a novel sodium-balanced chow (Envigo, TD.90229) (sodium-replete). Mice underwent the same procedure a second time 24h later and then were tested for behavior after an additional 24h. Mice were tested in the multi-spout assay under either sodium-deplete or sodium-replete states in a single session. Following 48h of ad-*libitum* access to water and standard laboratory chow, mice went through the furosemide treatment and behavioral testing again with the opposite homeostatic state.

### Fiber-Photometry

Wild-type mice underwent surgery for expression of dopamine sensors and fiber implantation as outlined above (see *Surgeries*) before undergoing multi-spout consumption of sucrose under different homeostatic demand states (see *Homeostatic Demand Multi-Spout Experiments*). We recorded dopamine dynamics in the NAc medial shell and lateral shell simultaneously during behavior in the multi-spout assay over 3 consecutive sessions in each homeostatic demand state. After head-fixation, we connected to the mouse’s fiber implant, patch cables (Doric, 400 µm, 0.37NA, 2.5 mm stainless steel ferrules) coupled to a 6-port mini cube (Doric, FMC6_IE(400-410)_E1(460-490)_F1(500-540)_E2(555- 570)_F2(580-680)_S) that was coupled to an integrated fiber-photometry system (Tucker-Davis Technologies, RZ10X). We delivered 405 nm and 465 nm light sinusoidal modulated at 211 Hz and 331 Hz, respectively. The average power for each wavelength was calibrated to 30 µW using a power meter (Lux integrated with the RZ10x) prior to the experiment. The fluorescent emission produced by 405 nm and 465 nm excitation were collected using the same fiber used to deliver light and were measured on a photodetector (Lux) and demodulated during recording. The timing of behavioral events were recorded via TTL communication to the fiber-photometry system.

Fiber-photometry was analyzed using custom Python and R scripts that are freely available through our GitHub (https://github.com/agordonfennell/open_stage). A custom Python script was used to convert raw data into tidy format and then an assortment of custom R functions were used to process the fiber-photometry signals. The decay in signal throughout the session for the 405 and 465 channels were corrected by fitting and subtracting a 3^rd^ degree polynomial to each raw signal. We then normalized the signals by computing *z*-scores using the mean and standard deviation of the entire session. Using the onset of each access period, we created perievent time histograms with time relative to access onset and then resampled signals to 20 samples per second. We used the 405 signal to assess movement artifacts but did not observe any abrupt changes in fluorescence that typically indicate such artifacts (**Figure 7- Figure Supplement 5**). The 405 signal was not used to correct the 465 signal because we observed simultaneous opposing signals in the 405 and 465 signal that may be attributed to the fact that 405 is not an ideal isosbestic signal for GRAB-DA. The 465 signal in perievent time histograms was shifted based on the mean signal during the 3 s prior to the onset of the access period. To summarize the dopamine signal during access to each solution, we computed the average and peak signal during the access period.

### Histology

We conducted *post hoc* histology to determine the location of viral expression and optical fiber locations for mice in optogenetic and fiber-photometry experiments. At the conclusion of the experiment, mice were deeply anesthetized and transcardially perfused with 20 mL 1x PBS and 20 mL 4% paraformaldehyde (PFA). Heads were removed and post-fixed in 4% PFA for 24h, brains were removed and post-fixed for an additional 24h, and then brains were transferred to 30% sucrose until they sank. Brains were frozen at -20°C and sectioned at 40 µm on a cryostat (Leica). Every other brain section was collected in 1x PBS and then mounted on a glass slide. Slides were cover-slipped using the mounting medium Fluoroshield with DAPI for visualizing cell nuclei. Sections that contained the bottom of the optic fiber were imaged with epifluorescence at 5x magnification (Zeiss, ApoTome2; Zen Blue Edition). The location of optic fibers was determined by mapping the position of the fiber using a mouse brain histology atlas (Paxinos and Franklin, 2001). The position of fibers was overlaid onto a vector image of the corresponding atlas section using Illustrator (Adobe).

### Statistical Analysis and Visualization

Results with repeated measures design were analyzed using a repeated measure analysis of variance (rmANOVA) using the afex package (0.28.1) in R. We computed *post hoc* comparisons using Tukey’s Honest Significant Difference (HSD) using the emmeans package (1.6.0) in R. Results with two variables were analyzed using a two-sided unpaired *t*-test using base R. Correlations were computed using Pearson’s product moment correlation coefficient using base R. For all statistics, significance was set at P values less than 0.05. Details for all statistical results presented in the paper can be found in the stats table included with this manuscript.

Data was visualized using the ggplot2 (3.3.3) package in R. Combined plots were assembled using patchwork (1.1.1). Color scales were produced using pals (1.7) or viridis (0.6.2). 3D renderings of the head-fixed hardware were produced using Fusion 360 (Autodesk). Plot components were assembled and further edited using Illustrator (Adobe).

## Acknowledgements

A.G.F was supported by F32 DA054719, T32 DA7278-27, R01 DA038168, R21 DA050868, and P30 DA048736. G.D.S was supported by R37 DA032750, R01 DA038168, R21 DA050868, and P30 DA048736. M.R.B. and R.G. were supported by R37 DA03339, P50MH119467 (project 4), and P30 DA048736. M.F.R. and P.B. were supported by R21 DA050868.

We would like to thank Dr. Vijay Namboodiri and Madelyn Hjort for providing initial training to A.G.F. in the Arduino language. Lydia Gordon-Fennell for creating vector illustrations of head-fixed mice in figure diagrams and editing the manuscript. Barbara Benowitz for editing the manuscript. Dr. Spencer Smith for providing initial designs for metal head-rings and head-fixation stage. Dr. Nick Steinmetz for providing insights into the behavior of mice during head-fixed behavior with wheel turning as an operant response.

**Figure 1- Figure Supplement 1:**
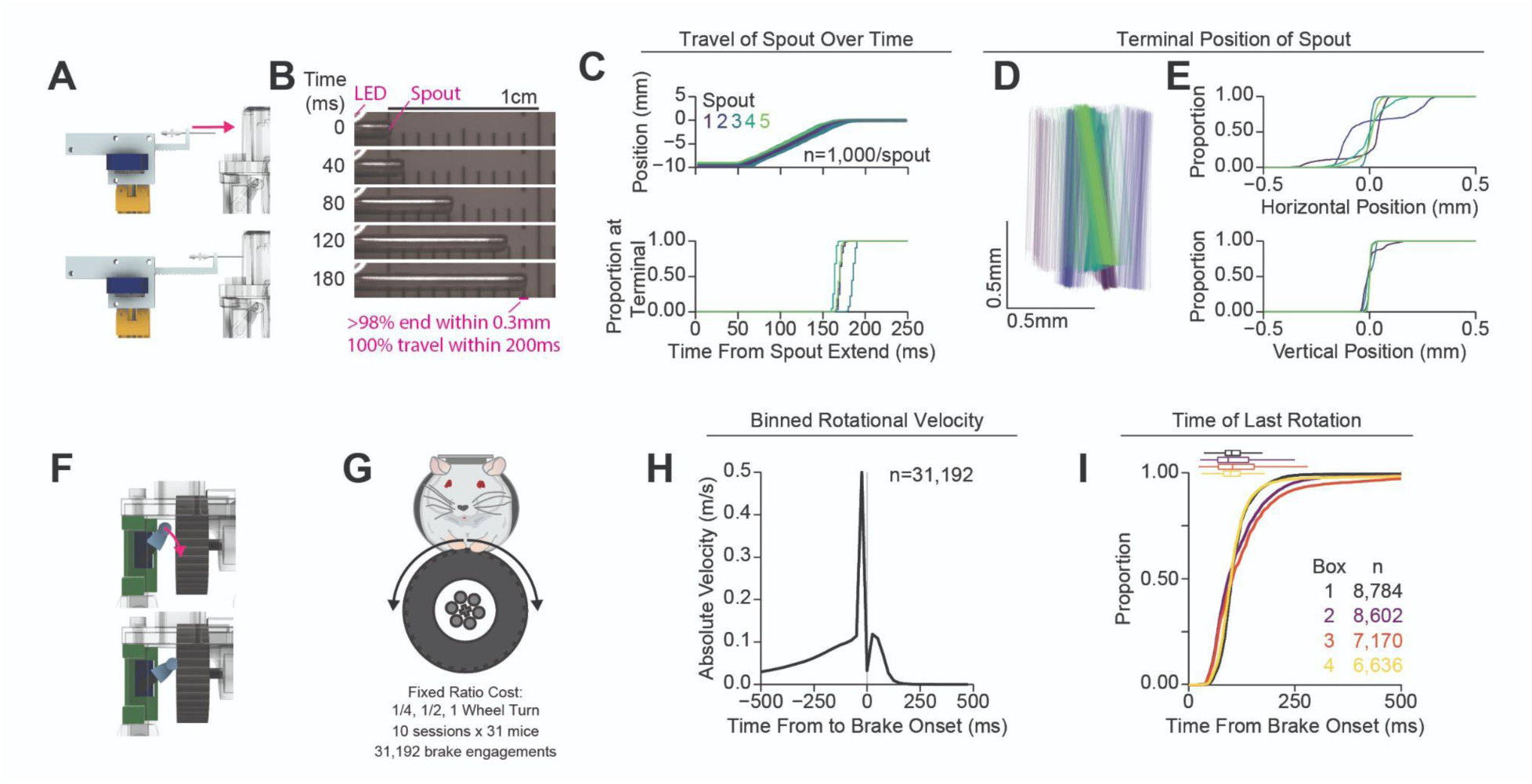
Validation of Retractable Spout and Wheel Brake. A) Cartoon of validation of the retractable spout. B) Representative still frames from video data recording the position of the spout during 1,000 extension/retractions in 5 different retractable spout assemblies. An LED was used to indicate the onset of the extension command (top left of frame). C) Horizontal position of the spout during extension (top) and cumulative distribution of latencies to reach 98% of the final spout position (bottom). D) Terminal position of the spout on all recorded extensions drawn from tracking the top and bottom corners of the spout (color indicates the identity of the spout assembly, lines represent the tip of the spout in 2D space for each trial). E) Cumulative distribution of the horizontal (top) and vertical (bottom) position of the spout at the terminal location relative to the mean position at the terminal location, indicating incredibly consistent vertical positioning of the spout and highly consistent horizontal positioning of the spout. F) Cartoon of validation of the wheel brake. G) Cartoon indicating that data from fixed-ratio self- administration of 10% sucrose was used for (H-I) (n=31,192 wheel brake engagements). H) Mean absolute velocity during 25ms bins (SEM is not visible behind the line). I) Cumulative distribution of the time of last detected rotation across every trial for each of the 4 behavioral setups (box and whisker plots indicate the median and interquartile range).

**Figure 1 - Figure Supplement 2:**
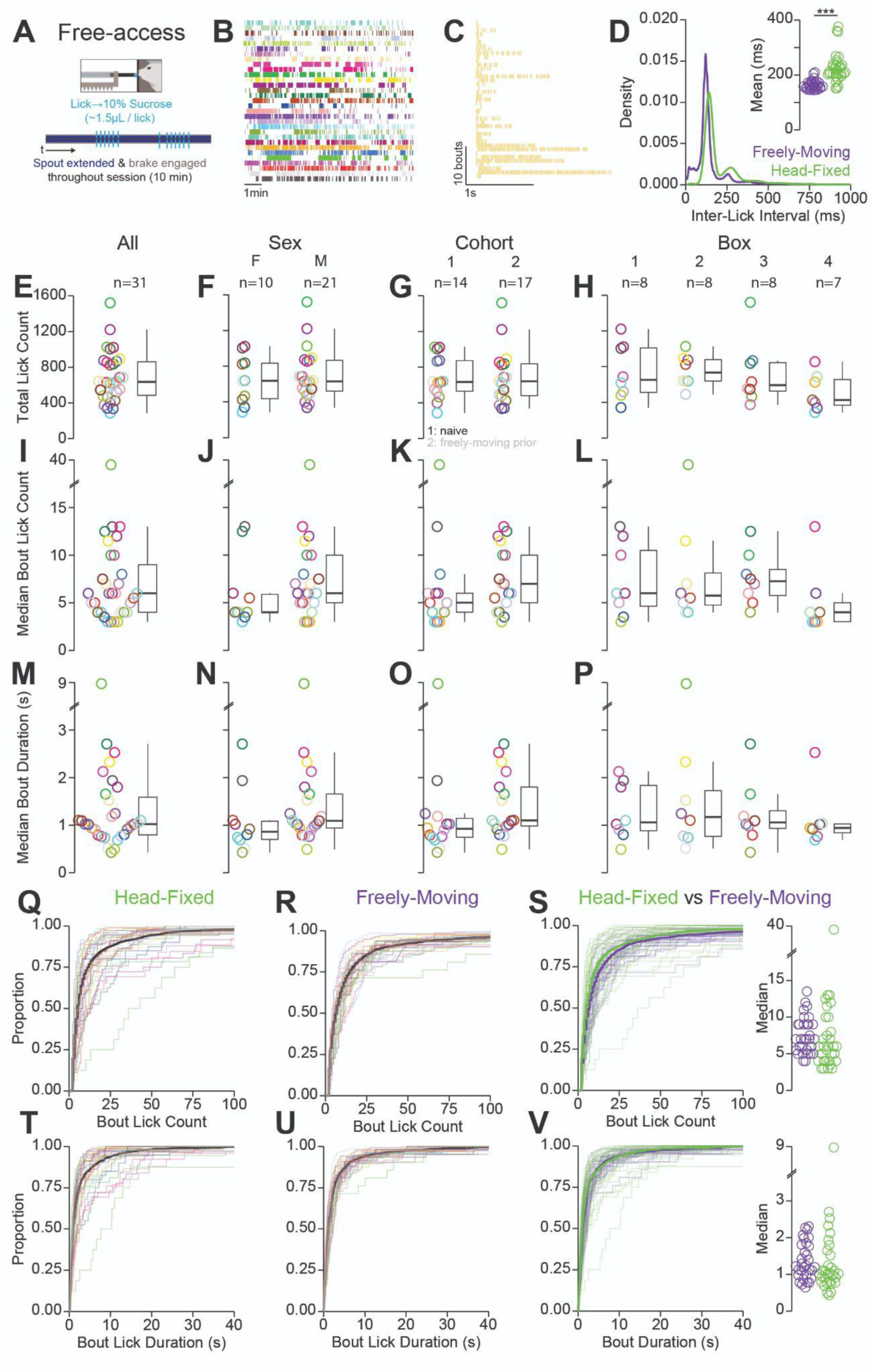
Quantification of Behavior During Head-Fixed Spout Training. A) Cartoon depicting the task design for head-fixed, free-access consumption of sucrose (free-access lick training). B) Raster of all licks recorded in a single 10 min session for the 31 mice in the experiment (color indicates the mouse identity and is consistent with other plots depicting color-coded single mouse data). C) Raster of all licking bouts for a representative mouse. D) Density plot of inter-lick intervals in the head-fixed (green) and freely moving (purple) versions of free-access consumption. Inset shows the median inter-lick interval for all 31 mice in both versions of the task (*t*-test: freely moving vs. head-fixed***). (E-P) Summaries of licking behavior during free-access in the head-fixed version of the task depicting the total lick count (E-H), median lick count per licking bout (I-L), and median duration per licking bout (M-P). Each row contains the same data from 31 mice divided by sex, cohort (1: naïve, 2: freely moving prior), and behavioral box number. No statistically significant differences in free-access licking behavior were observed. (Q-S) Cumulative distribution of the number of licks per bout in the head-fixed (Q) and freely moving (R) versions of the task. S) Data from (Q) and (R) overlayed and color coded based on the version of the task (left) and the median values for each mouse in both versions of the task (right). (T-V) Same as (Q-S) but for the duration of licking bouts (*rings and faded lines depict individual mice; no statistical differences for any comparison; see stats table for details*).

**Figure 1 - Figure Supplement 3:**
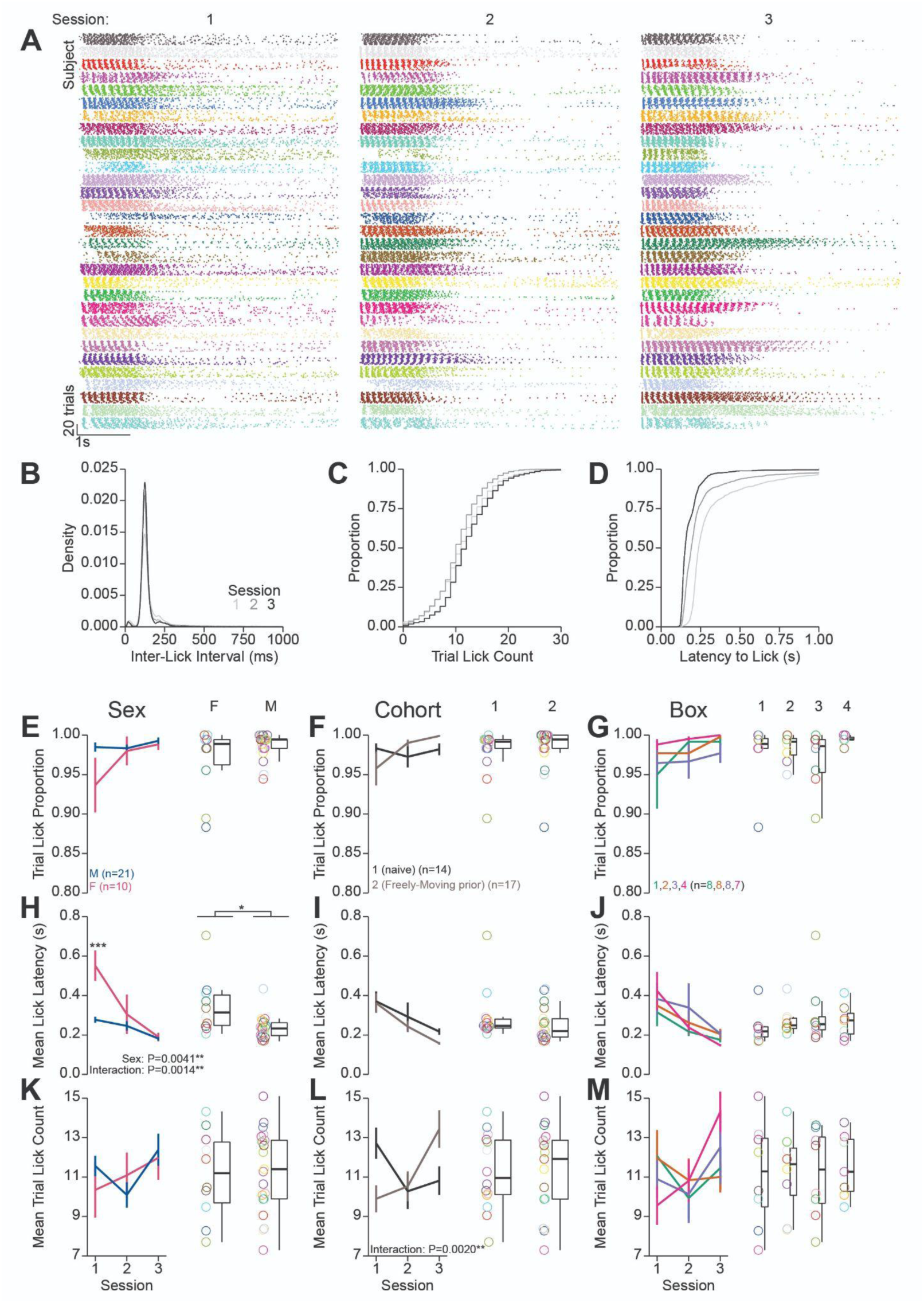
Quantification of Behavior During Head-Fixed Retractable Spout Training. Further quantificatino of behavior during retractable spout training corresponding to data shown in Figure 1G-J. A) Lick raster of all licks recorded across all trials across 3 sessions for the 31 mice in the experiment (color indicates the mouse identity and is consistent with other plots depicting color coded single mouse data). B) Density of inter-lick intervals. C) Cumulative distributions of number of licks per trial. D) Cumulative distribution of latency from spout extension command to first lick on trials with at least one lick. (E-M) Summaries of behavior during retractable spout training in each session (left) and averaged across sessions (right) depicting the proportion of trials with a lick (D-G), mean latency from spout extension to first lick (H-J), and mean number of licks per trial (K-M). Each row contains the same data from 31 mice divided by sex, cohort, and behavioral box. (*Color in (B-D) represents the session number; rings depict individual mice; effects listed on plots indicate statistical significance for significant Two-Way RM ANOVA effects; asterisks above means indicate significant differences determined with HSD across group within the same session; asterisks above box and whisker plots indicate significant differences determined with a t-test or One-Way ANOVA; see stats table for details*).

**Figure 1 - Figure Supplement 4:**
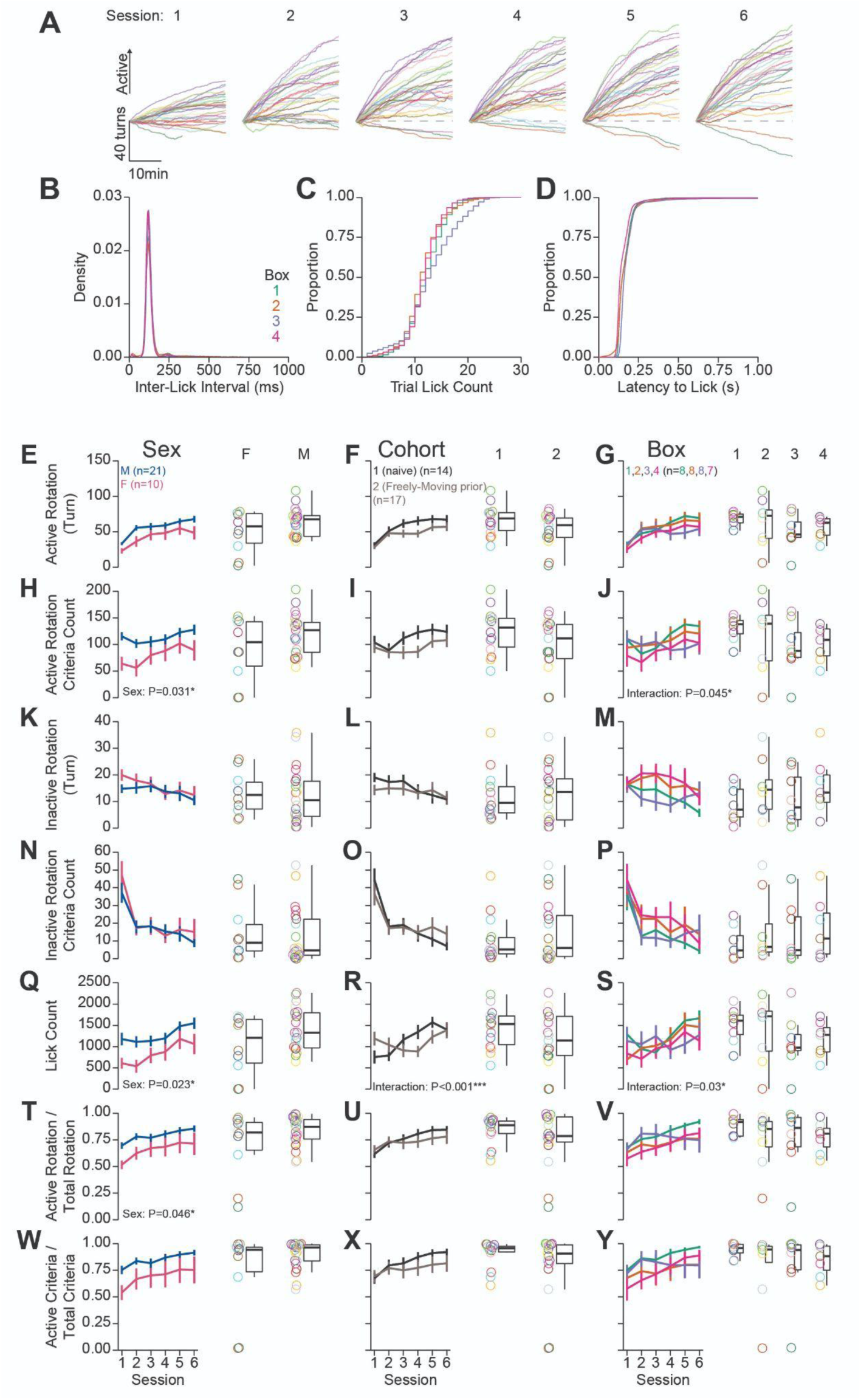
Quantification of Behavior During Head-Fixed Operant Conditioning for Sucrose. Further quantification of behavior during operant training corresponding to data shown in Figure 1L-N. A) Cumulative position of the wheel for all mice across all sessions. B) Density of inter-lick intervals. C) Cumulative distributions of number of licks per trial. D) Cumulative distributions of latency from spout extension to first lick. (E-Y) Summaries of behavior during operant training depicting numerous metrics across each row. Each row contains the same data from 31 mice divided by sex, cohort, and behavioral box number. Line graphs depict the mean and SEM for each session of training, the box and whisker plots with individual mice plotted as rings depicts the *mean over the last 3 sessions*. (*Color in (B-D) represents the box number; rings and faded lines depict individual mice; Effects listed on plots indicate statistical significance for significant Two-Way RM ANOVA effects; asterisks above means indicate significant differences determined with HSD across group within the same session; asterisks above box and whisker plots indicate significant differences determined with a t-test or One-Way ANOVA; see stats table for details*).

**Figure 1 - Figure Supplement 5:**
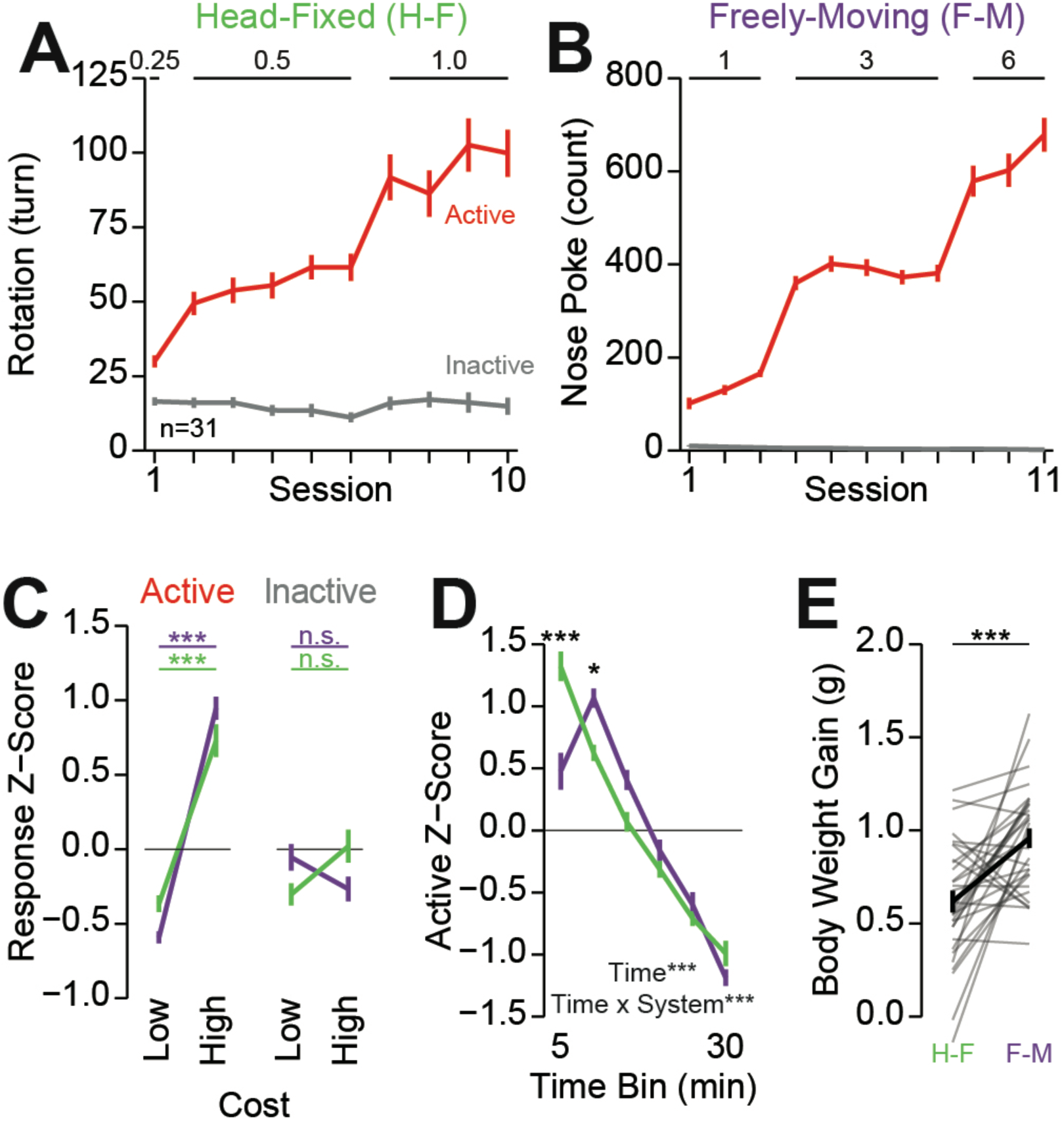
Comparison of Head-Fixed and Freely moving Versions of Operant Conditioning. A) The total rotation of the wheel across all sessions of operant conditioning in the head-fixed version of the task (numbers on the top of the plot indicate the cost of reward in wheel turns; data in the last 3 sessions of 0.5 and 1.0 correspond to data presented in Figure 1N). B) Total nose poke count across all sessions of operant conditioning in the freely moving version of the task (numbers on top of the plot indicate the cost of reward in nose pokes). C) Mean *z*-score across the last 3 sessions of responding during the low (0.5 turn/3 nose pokes) and high (1.0 turn/5 nose pokes) cost sessions (color of the line indicates the system; horizontal lines indicate HSD comparisons within a response; no significant difference between systems at any cost for a particular response (e.g. head-fixed active low vs freely moving active low)). D) Mean *z*- score of the active response during 5 min bins within the last 3 sessions of high cost sessions (asterisk above means indicates significant HSD comparison across systems at a given time bin). E) Total body weight gain (liquid consumption) during the head-fixed and freely moving versions of the task (*t*-test: Head-fixed vs. freely moving***). (*See stats table for details*).

**Figure 1 - Figure Supplement 6:** Video of Operant Responding for 10% Sucrose. Video of operant responding for 10% sucrose under a fixed-ratio (FR) of 1/2 turn.

**Figure 2 - Figure Supplement 1:**
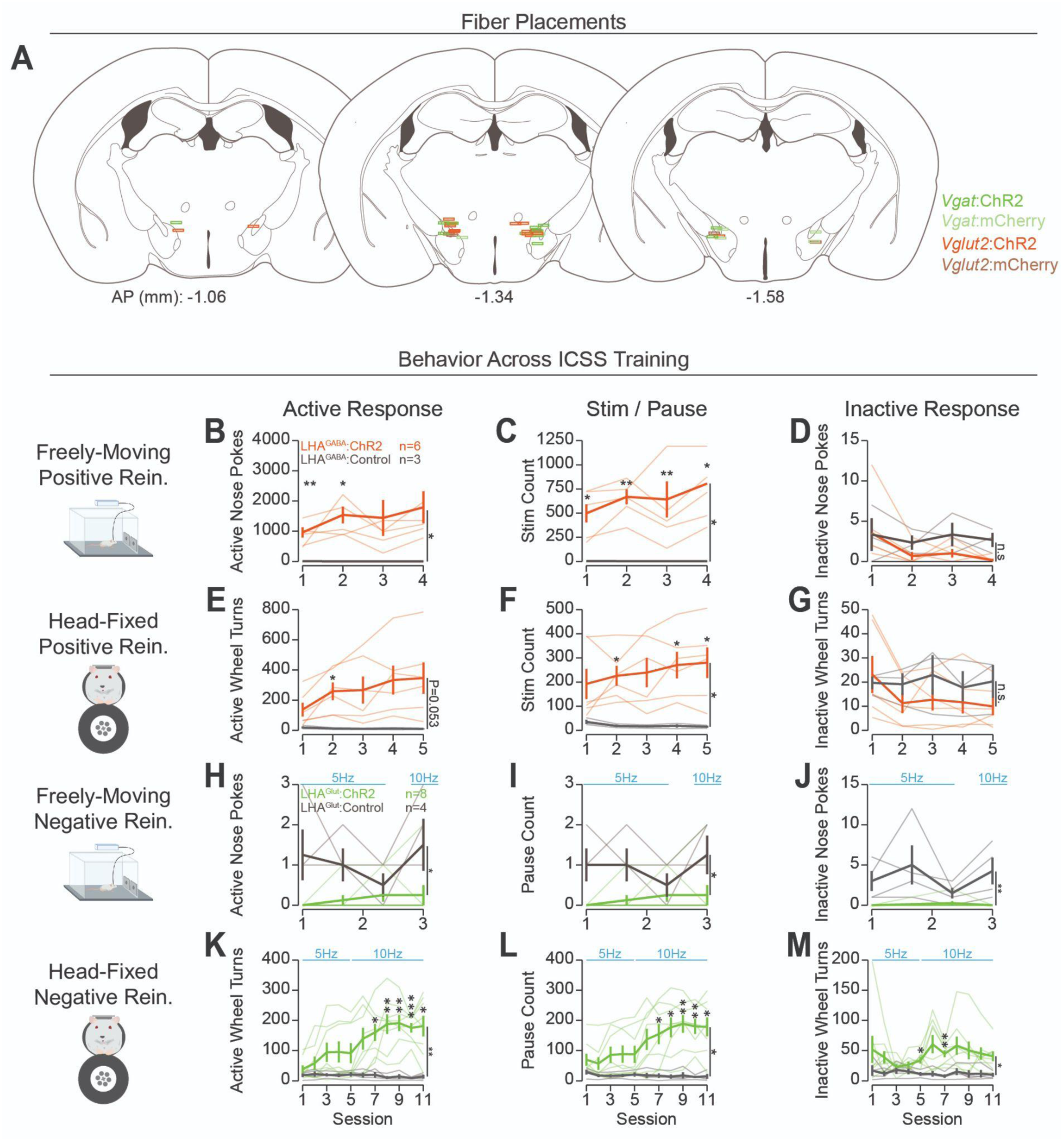
Placement of Optic Fibers and Training Data for Operant Conditioning to Obtain or Avoid Optogenetic Stimulation. A) Histological locations relative to bregma of the tip of optic fibers targeting the LHA. Colors indicate the experimental group of the corresponding mouse. (B-M) Training data across training for each stage of the task with counts for active responses (left column), stimulations or pauses (mid column), and inactive responses (right column). (B-D) Behavior during freely moving positive reinforcement. (E-G) Behavior during head-fixed positive reinforcement. (H-J) Behavior during freely moving negative reinforcement. (K-M) Behavior during head-fixed negative reinforcement. (*Asterisks depict Two-Way RM ANOVA main effects (vertical lines) or Bonferroni adjusted t-test comparisons between a group at a corresponding session (over means); Faded lines depict individual mice; see stats table for details*).

**Figure 2 - Figure Supplement 2:** Video of Head-fixed Operant Conditioning to Obtain Stimulation of LHAGABA Neurons or Avoid Stimulation of LHAGlut Neurons. Videos showing responding for optogenetic stimulation of LHA^GABA^ neurons under a positive reinforcement schedule (left) and responding for optogenetic stimulation of LHA^Glut^ neurons under a negative reinforcement schedule (right). The LED near the center of the frame indicates when the optogenetic stimulation is turned on under positive reinforcement or when the optogenetic stimulation is turned off under negative reinforcement.

**Figure 3 - Figure Supplement 1:**
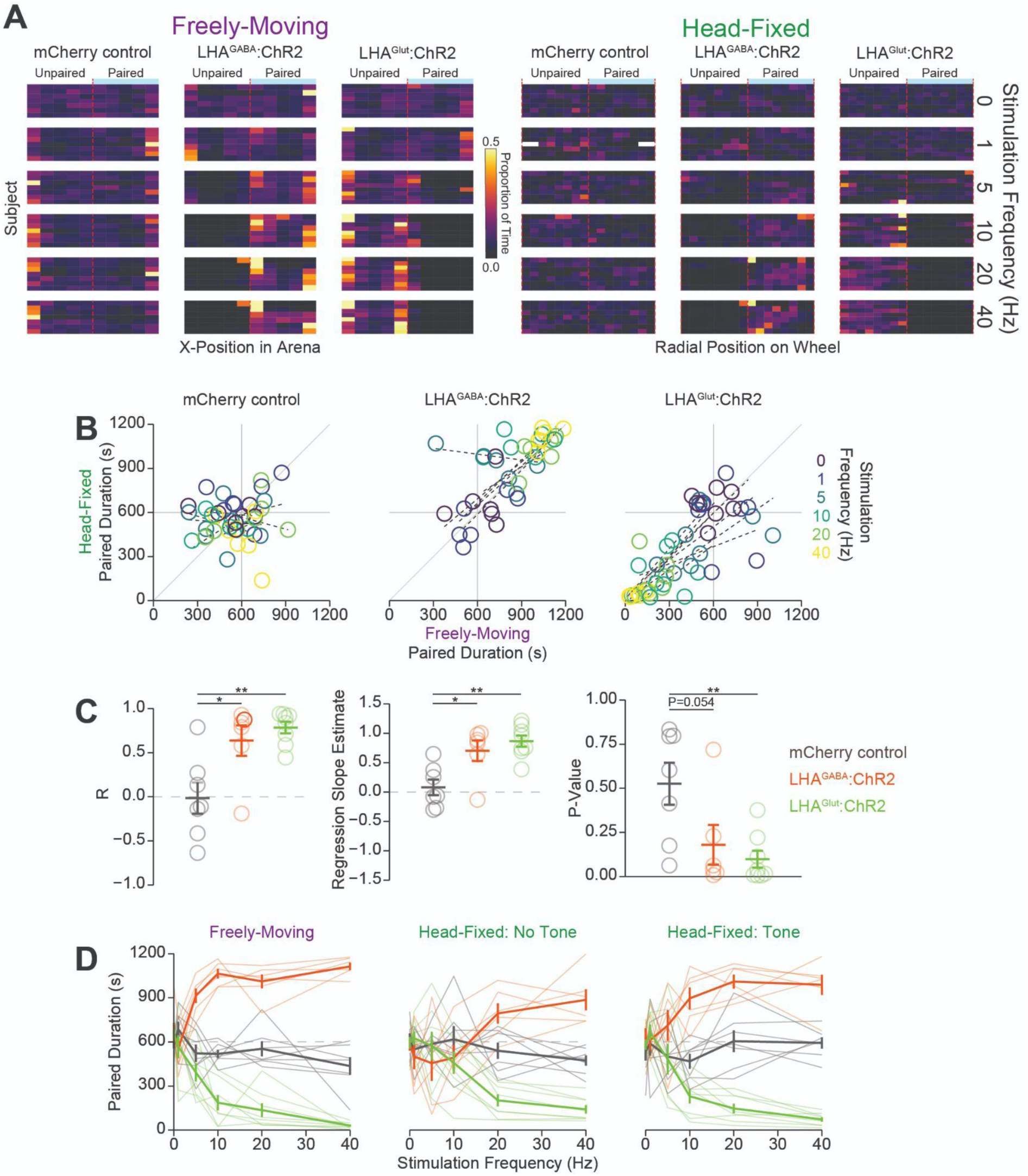
Quantification of Head-fixed and Freely moving Real-Time Place Testing. A) Heat map showing the binned x-position in the freely moving version of the task (left 3 columns) or radial position in the head- fixed version of the task (right 3 columns) of all mice across all stimulation frequencies (indicated by label on right). Color represents the proportion of time spent in the binned position. The paired side was counterbalanced across mice and sessions and is set to the right side for display purposes only. B) Correlation between the time spent in the paired zone during the freely moving (abscissa) or the head- fixed (ordinate) at different stimulation frequencies represented as different colors for individual mice. C) Summary statistics of correlations shown in (B) showing: R (left), estimated slope (middle), and P-value (right). D) Amount of time spent in the paired zone across the freely moving (left), head-fixed without a tone indicating the area the mouse was located in (middle), and head-fixed with a tone indicating the area the mouse was located in (right). (*Colors in (C-D) indicate the experimental group as indicated by the colors in the right side of C*; *asterisks depict HSD comparisons between groups as indicated by horizontal lines*).

**Figure 4 - Figure Supplement 1:**
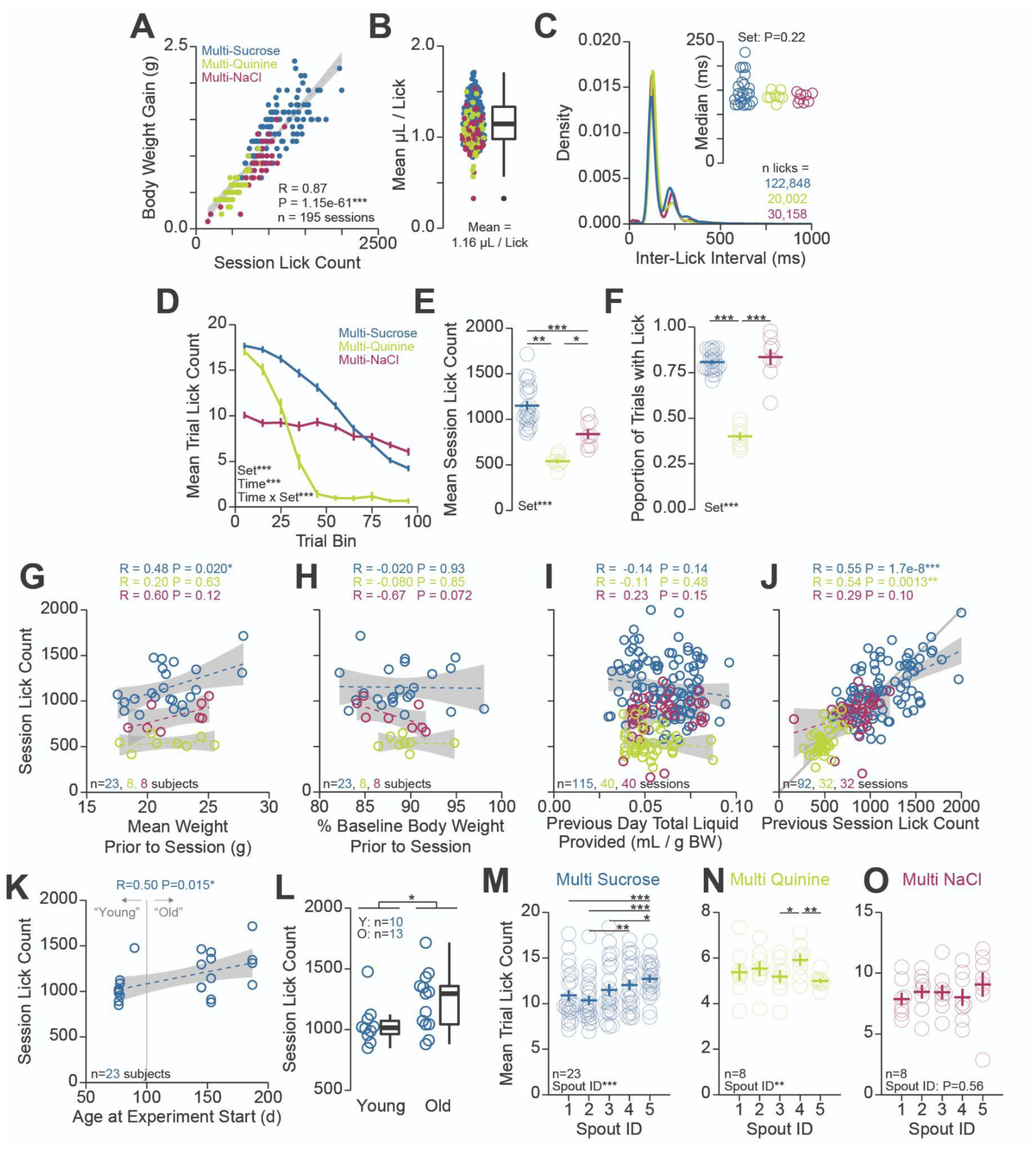
Quantification of Head-Fixed Consumption During Brief Access. A) Correlation between the number of licks and body weight gain within each session. B) Estimated volume consumed per lick produced by dividing the body weight gain by the number of licks within the session. C) Density plot of inter-lick intervals within each session set. Inset shows the median inter-lick interval. D) Mean trial lick count across all concentrations in bins of 10 trials across the session for each solution set. E) Mean total number of licks for sessions for each solution set. F) Proportion of trials with at least 1 lick for each solution set. (G-J) Correlations between different factors (abscissa) and session lick count (ordinate): G) mean weight prior to the session, H) percent baseline body weight prior to the session, I) total liquid consumed + provided to the mouse on the session prior to the session, J) the lick count on the previous session. K) Correlation between the age of the mouse at the experiment start (abscissa) and session lick count (ordinate) during the multi-spout consumption of a gradient of sucrose concentrations (mice used in multi- spout consumption of a gradient of NaCl concentrations and quinine concentrations did not have enough variance in age to conduct this analysis). L) Mean session lick count in mice defined as young (<100d at start of experiment) and old (>100d at start of experiment) (*t*-test: Young vs. Old*). M) Mean trial lick count across spouts for data presented in Figure 4. Although there were spout effects observed, counterbalancing solution and spout pairings across sessions controls for these effects. (*Main effects listed on plots are results of One-Way ANOVA (C-F) or One-Way RM ANOVA (M-O); asterisks depict HSD comparisons between solution sets indicated by horizontal lines; Faded lines and rings depict individual mice; see stats table for details)*.

**Figure 4 - Figure Supplement 2:**
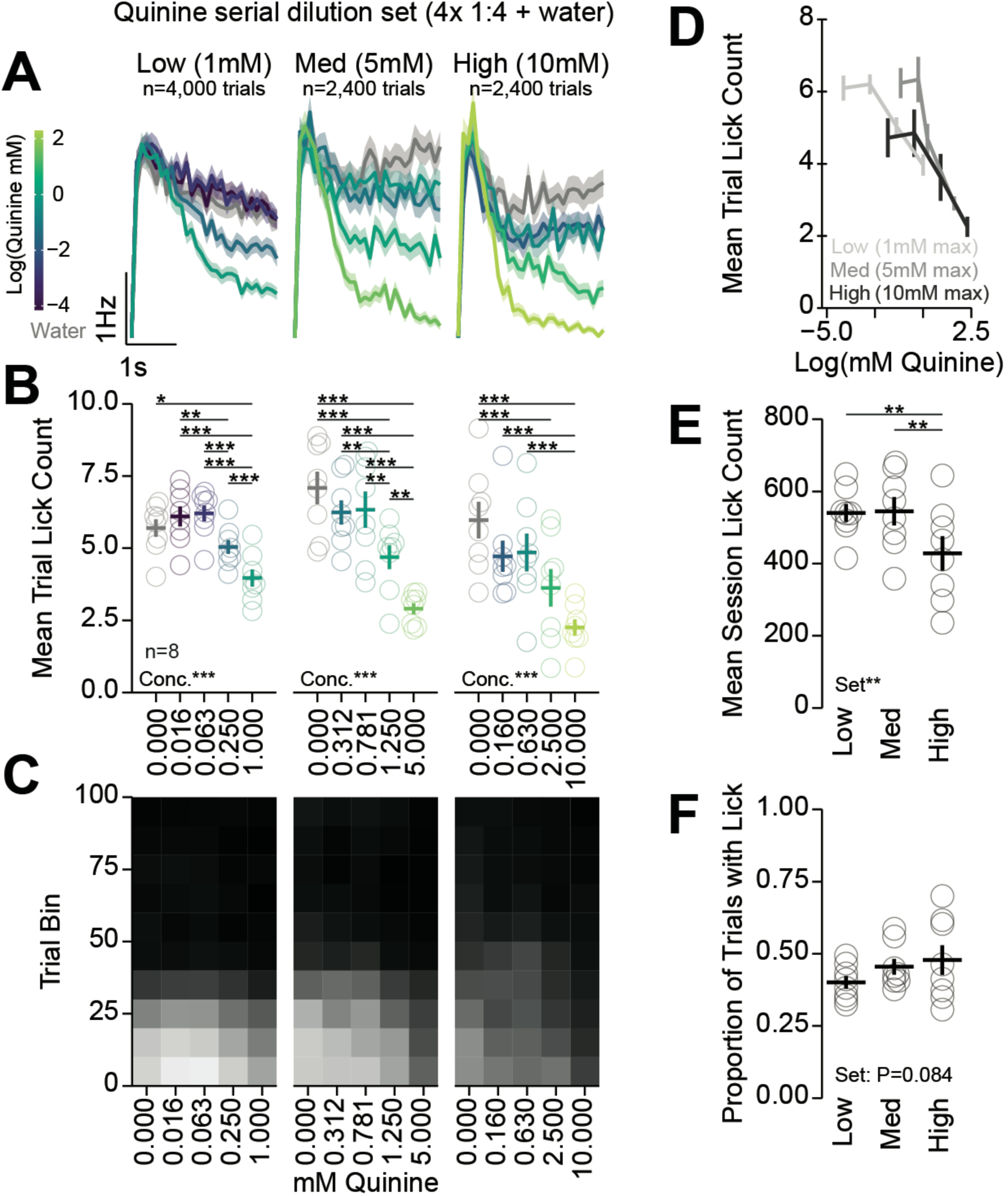
Multi-Spout Consumption of Different Gradients of Concentrations of Quinine. Analysis of 3 gradients of concentrations of quinine. Each concentration set had water and 4 concentrations of quinine with a 1:4 serial dilution starting at 1 mM (low), 5 mM (med), and 10 mM (high). (A-C) left, middle, and right columns depict data from low, med, and high concentration sets of quinine. A) Mean binned lick rate for all mice for each concentration. B) The mean number of licks per trial for each concentration. C) The mean number of licks for each concentration per trial binned by 10 trials over the course of the session. D) The mean number of licks per trial for quinine concentrations greater than 0 shown on a log scale of quinine concentration on the abscissa. E) Mean session lick count for each concentration set. F) Proportion of trials with a lick for each concentration set. (*Main effects listed on plots are results of One- Way RM ANOVA; asterisks depict HSD comparisons indicated by horizontal lines; Faded rings depict individual mice; see stats table for details)*.

**Figure 4 - Figure Supplement 3:**
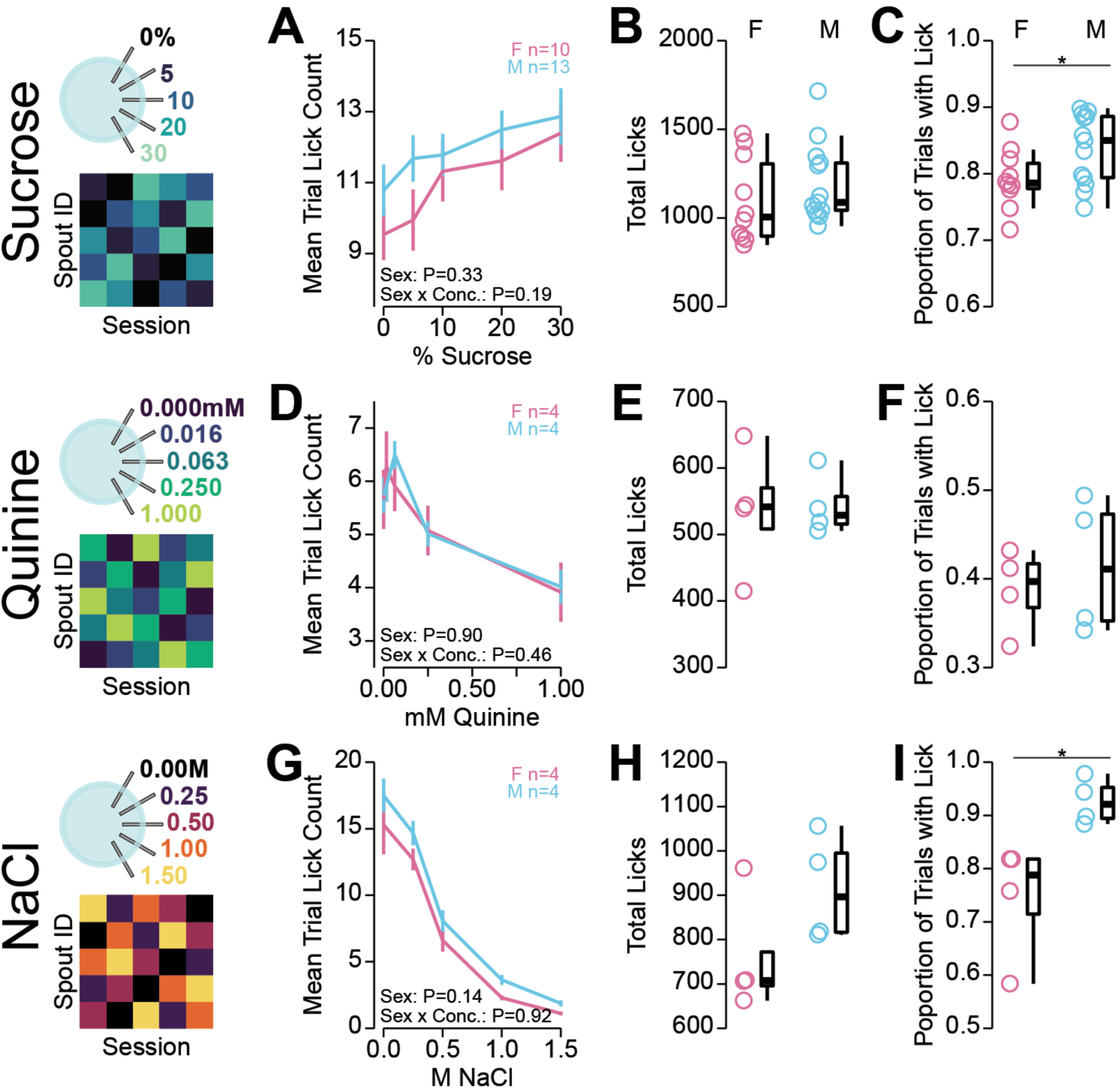
Sex differences in Head-Fixed Multi-Spout Consumption Behavior. Investigation of potential sex effects on behavior in the multi-spout brief-access assay shown in Figure 4. (A-B) Consumption of a gradient of concentrations of sucrose. A) The mean number of licks per trial for each concentration. B) Mean session lick count. C) Proportion of trials with a lick. (D-F) same as (A-B), but for consumption of a gradient of concentrations of quinine. (G-I) same as (A-B), but for consumption of a gradient of concentrations of NaCl. (*Main effects listed on plots are results of Two-Way RM ANOVA; no differences between sexes at a corresponding concentration; asterisks indicate sex differences determined by t-test; see stats table for details)*.

**Figure 4 - Figure Supplement 3:** Video of Consumption Behavior in the Multi-NaCl Assay Under Water-Restriction. Video shows licking behavior during the first 25 trials of the multi-spout assay for gradients of NaCl concentrations under water-restriction. Each video depicts a single 3 s trial played back at half-speed. Videos are organized to display trials from top to bottom (earlier trials on the top), and NaCl concentration from left to right (lower concentrations on the left). However, concentrations were provided in pseudorandom order.

**Figure 5 - Figure Supplement 1:**
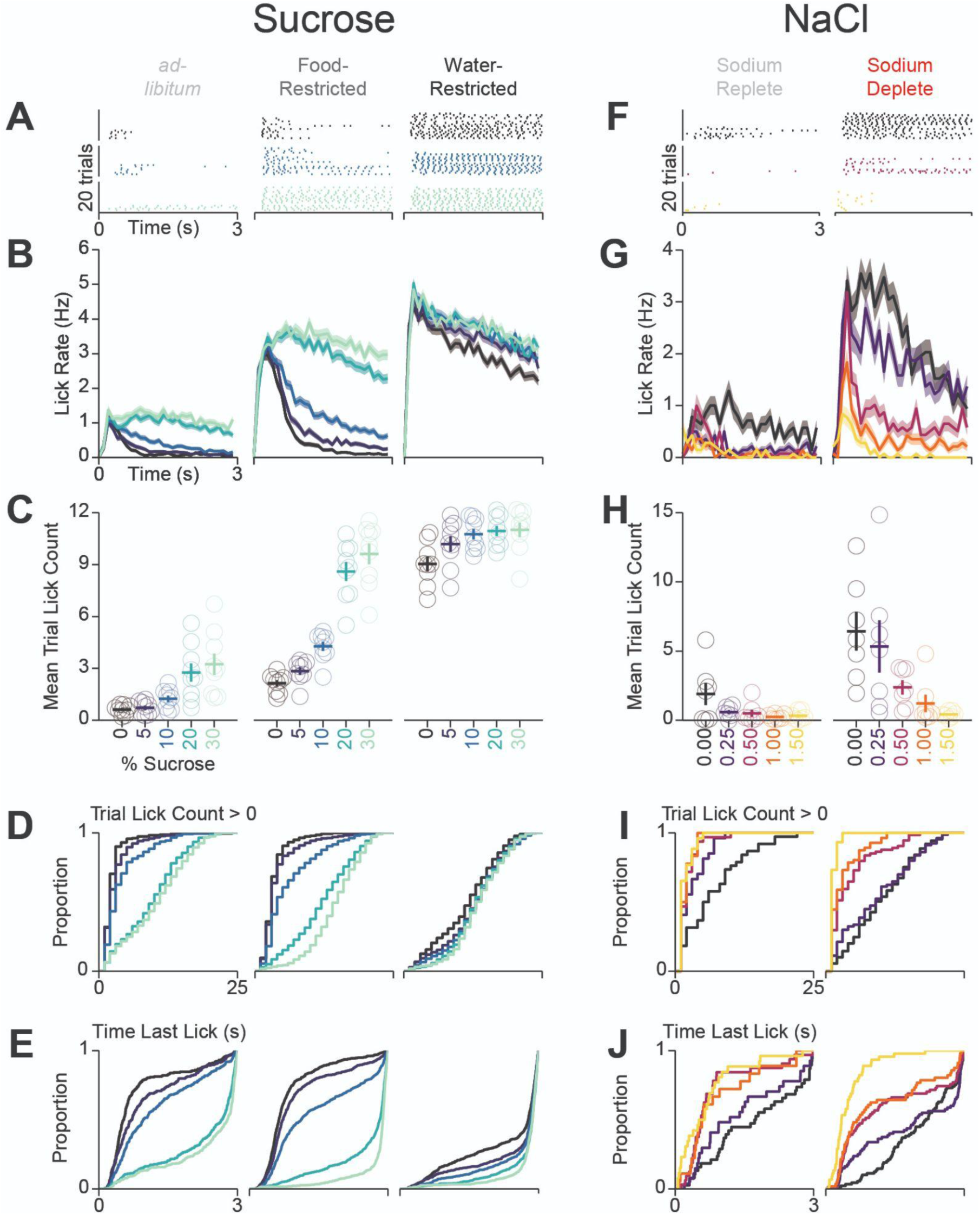
Behavioral Details for Differences in Consumption Across Homeostatic Demand. : Sets of columns containing data from mice in Figure 5 undergoing multi-spout consumption of a gradient of concentrations of sucrose (left 3 columns) or NaCl (right 2 columns) across homeostatic demands. A) Lick raster of a representative mouse depicting the licks for water, medium concentration, and high concentration during the 3 s access period. B) Mean binned lick rate for all mice for each concentration. C) The mean number of licks per trial for each concentration (same data as Figure 5 but with single mice displayed). (D-E) Cumulative distribution of the number of licks in trials with a lick (D) and the time of the last lick within each licking bout (E). (F-J) same as (A-E), but for consumption of a gradient of concentrations of NaCl.

**Figure 6 - Figure Supplement 1:**
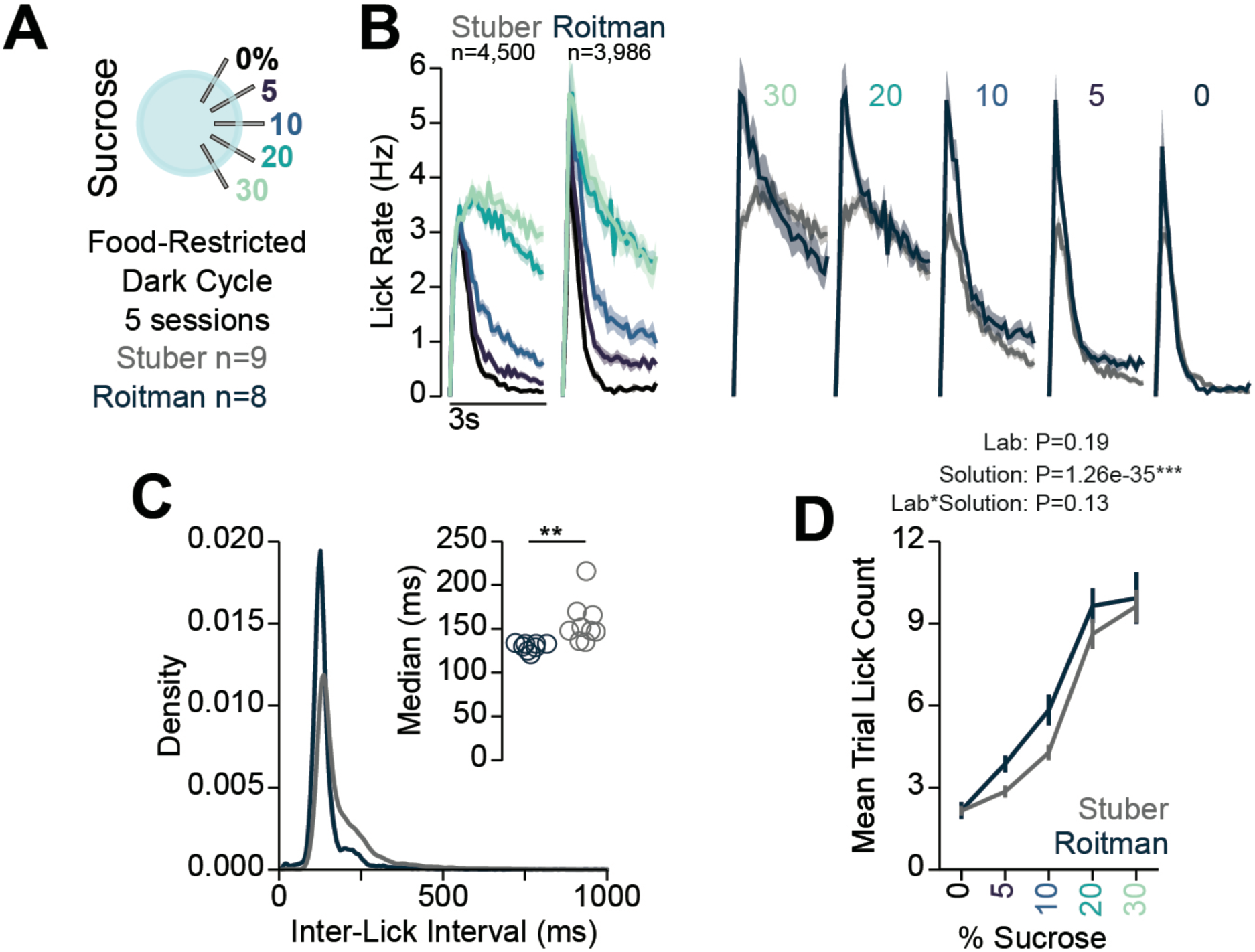
Comparison of Multi-Spout Behavior Across Labs. : A) Data included in figure: comparison between mice that were food-restricted and ran through multi-spout brief-access to a gradient of sucrose concentrations in the dark-cycle in the Stuber lab (data shown in Figure 5) and the Roitman lab (data shown in Figure 6). B) Mean binned lick rate for all mice for each concentration indicated by color with facets for lab (left) and for each lab indicated by color with facets for sucrose concentration (right). C) Density plot of inter-lick intervals for mice ran in the Stuber and Roitman labs indicating a higher density of low inter-lick-intervals in mice ran in the Roitman lab compared to the Stuber lab. Inset shows the median inter-lick interval (*t*-test**). D) The mean number of licks per trial for each concentration of sucrose for mice ran in the Stuber and Roitman labs indicating no effect of lab (Two-Way RM ANOVA stats listed on plot).

**Figure 7 - Figure Supplement 1:**
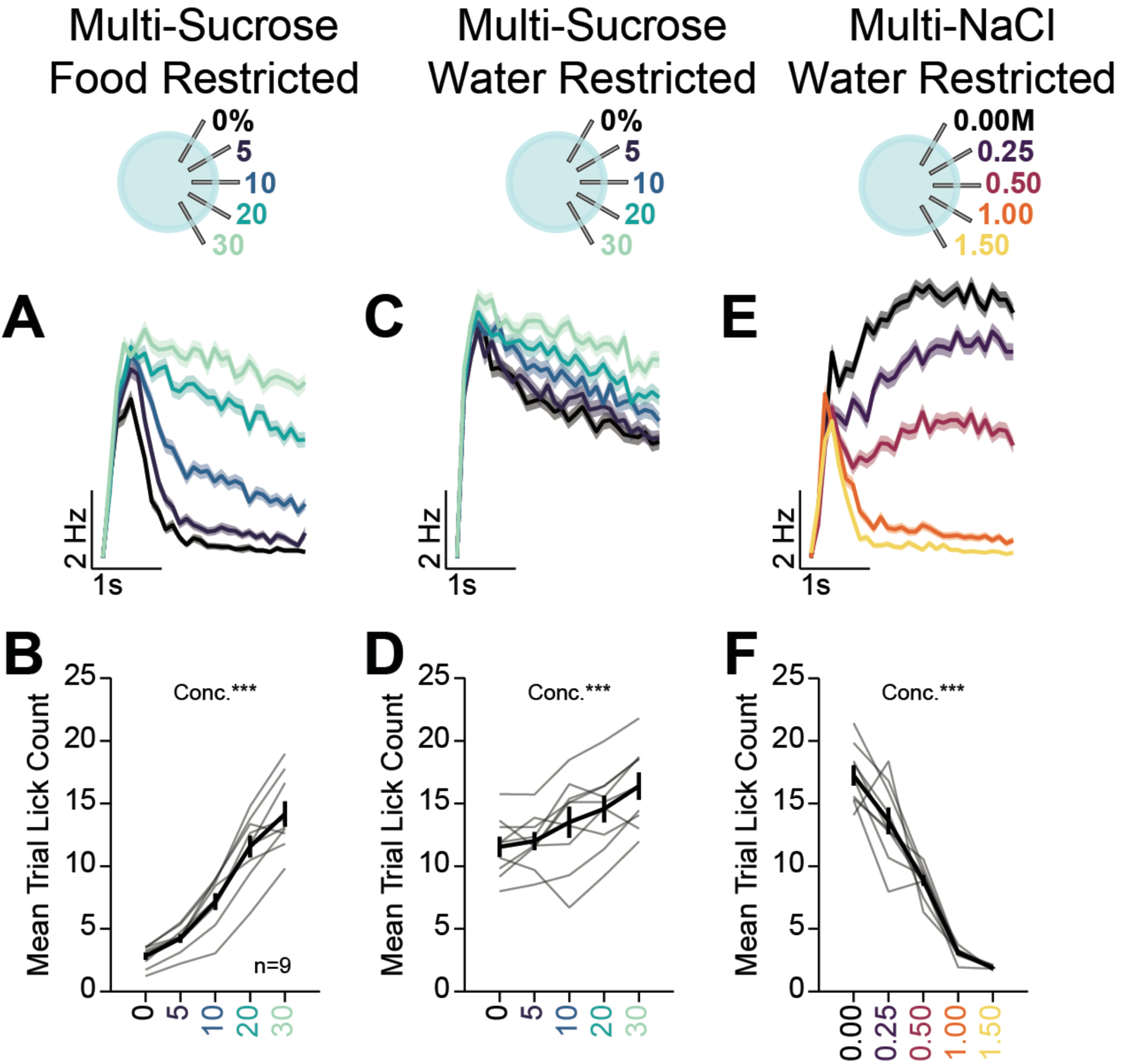
Multi-Spout Licking Behavior. Multi-spout licking behavior corresponding to Figure 7. A) Mean binned lick rate for all mice for each concentration during multi-spout consumption of sucrose under food-restriction (color indicates concentration of sucrose). B) The mean number of licks per trial for each concentration of sucrose under food-restriction. C) Mean binned lick rate for all mice for each concentration during multi-spout consumption of sucrose under water-restriction (color indicates concentration of sucrose). D) The mean number of licks per trial for each concentration of sucrose under water-restriction. E) Mean binned lick rate for all mice for each concentration during multi-spout consumption of NaCl under water-restriction (color indicates concentration of NaCl). F) The mean number of licks per trial for each concentration of NaCl under water-restriction (*Asterisks listed on plots are the results from One-Way RM ANOVA; see stats table for details*).

**Figure 7 - Figure Supplement 2:**
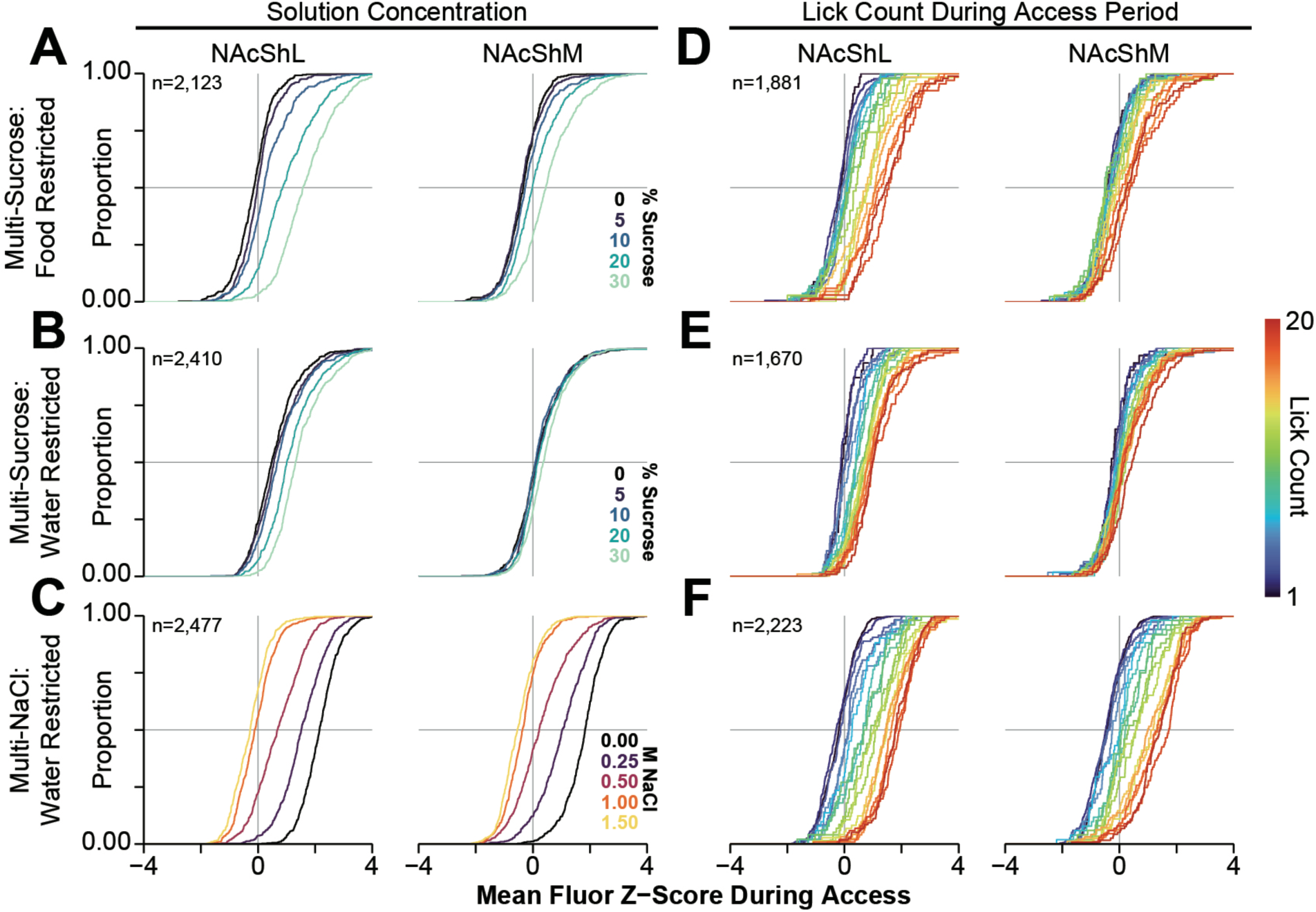
Cumulative Distribution Functions of GRAB-DA Responses in the NAcSh During Multi-Spout Consumption Behavior. (A-C) Cumulative distribution functions of mean GRAB-DA fluorescence during the access period for all trials with at least 1 lick during multi-spout sucrose under food-restriction (A), multi-spout sucrose under water-restriction (B), and multi-spout NaCl under water-restriction (C) (color indicates the solution ID as indicated in right inset). (D-F) Cumulative distribution functions of mean GRAB-DA fluorescence during the access period for all trials with 1-20 licks during multi-spout sucrose under food-restriction (D), multi-spout sucrose under water-restriction (E), and multi-spout NaCl under water-restriction (F) (color indicates the number of licks during the trial as indicated in right inset).

**Figure 7 - Figure Supplement 3:**
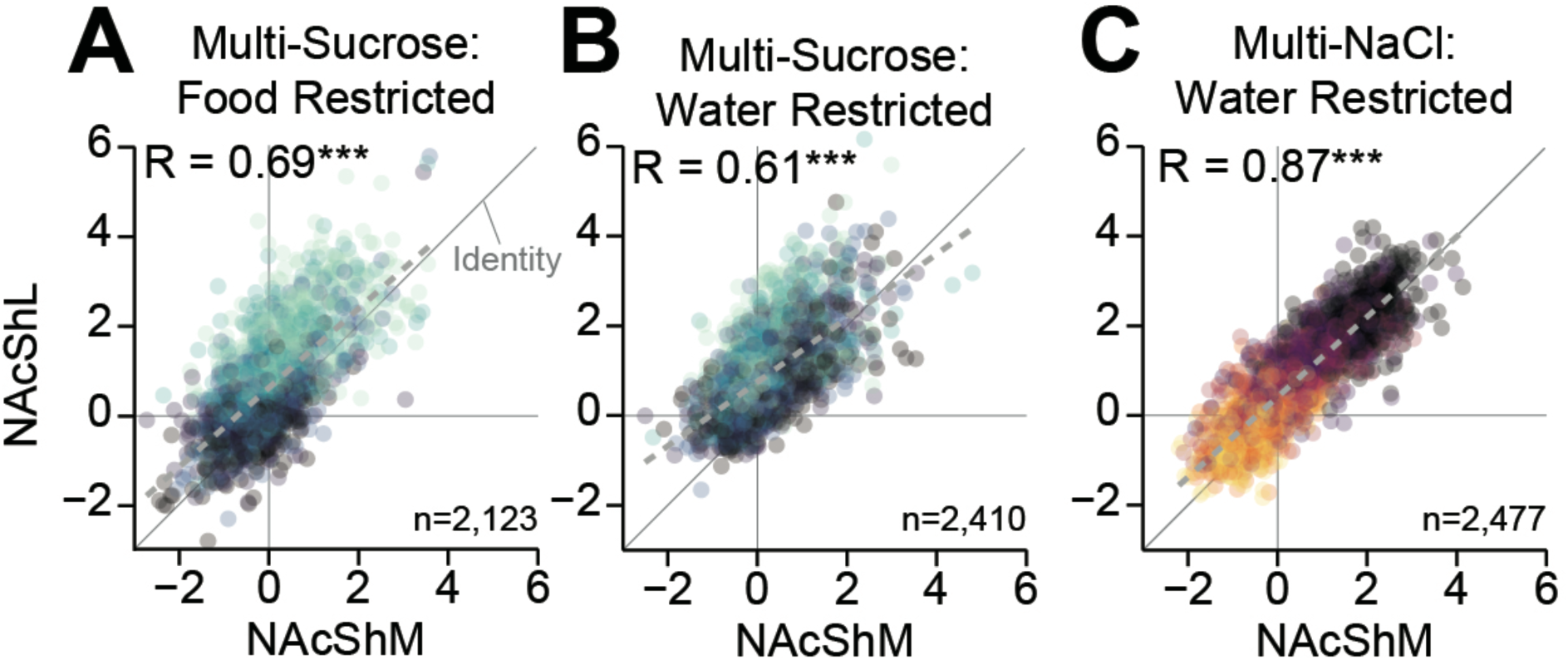
Linear Correlation of Dopamine Dynamics during Multi-Spout Consumption. (A-C) Correlations between NAcShM and NAcShL mean GRAB-DA fluorescence during the access period for all trials with at least 1 lick during multi-sucrose under food-restriction (A), multi-sucrose under water-restriction (B), and multi-NaCl under water-restriction (C) (\*\*\**Correlation P<0.001; see stats table for details*).

**Figure 7 - Figure Supplement 4:**
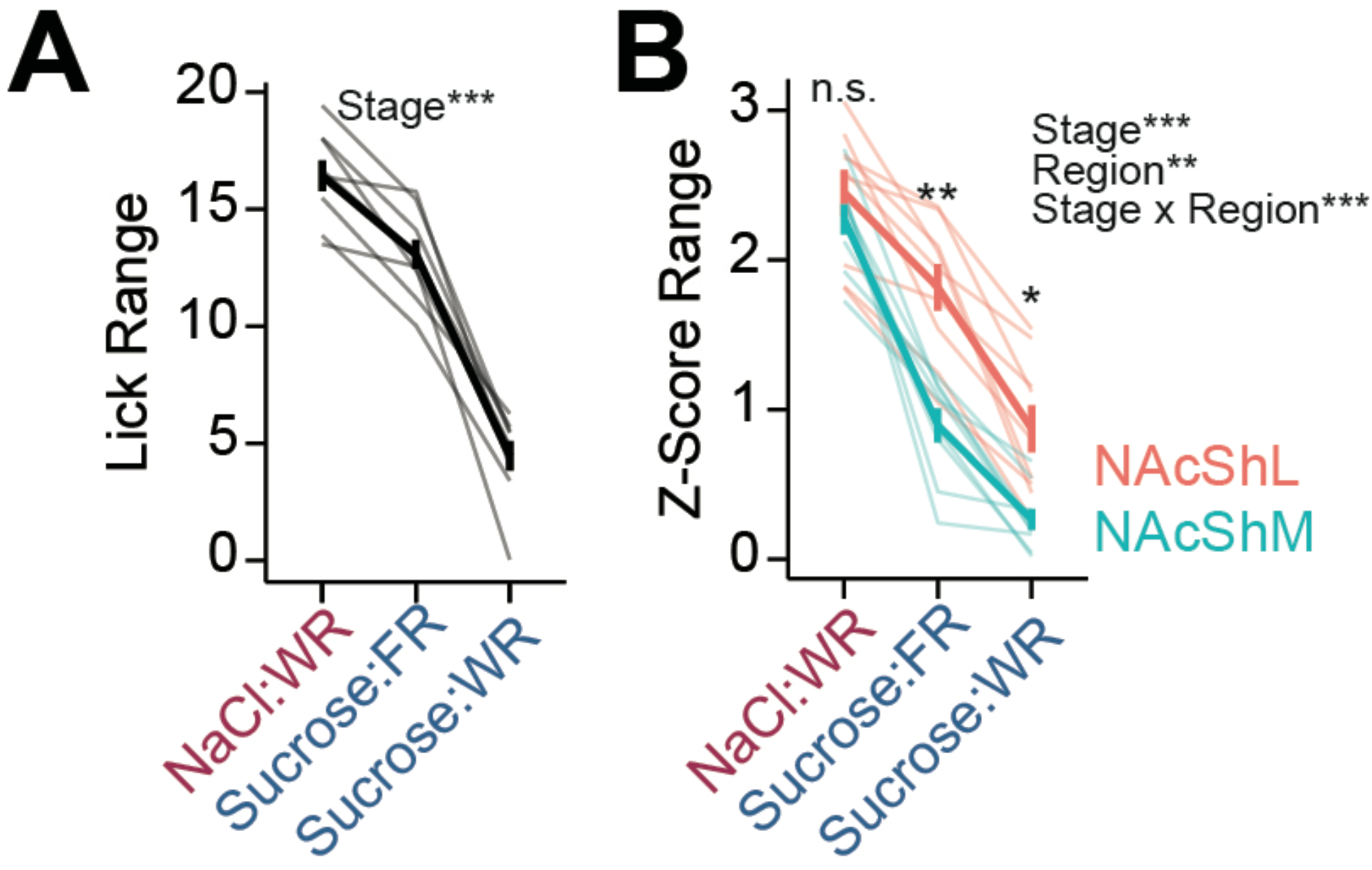
Range of Licking and NAcSh Dopamine Signals during Multi-Spout Consumption Behavior. (A) Range of licking (absolute difference in licking during access to highest and lowest concentrations) across each stage of the task (WR: water-restricted, FR: food-restricted) (*One-Way RM ANOVA: set\*\*\**). (B) Range of GRAB-DA fluorescence signals (absolute difference in mean *z*-score during access to highest and lowest concentrations) across each stage of the task (*Stats listed on plots are the results from One- Way RM ANOVA; asterisks indicate differences between NAcShM and NAcShL at a corresponding set; all comparisons across stages within a brain region are significant; see stats table for details*).

**Figure 7 - Figure Supplement 5:**
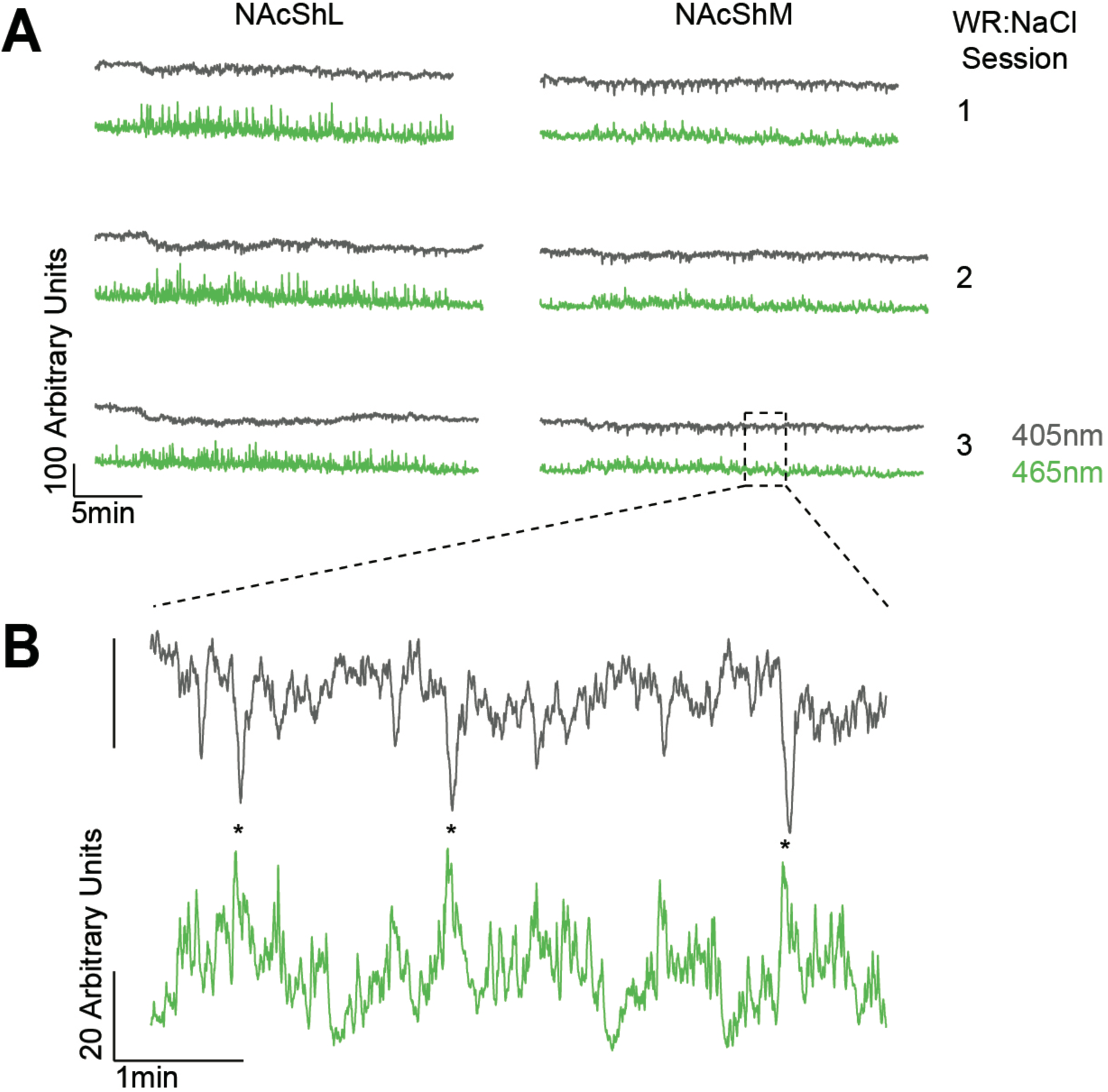
Representative Full-Session Traces. (A) Full-session raw traces from mouse abb11 across three sessions of multi-NaCl under water-restricted conditions (WR:NaCl). Both the 465 nm channel (used for GRAB-DA2m imaging) and the 405 nm channel (imprecise isosbestic) show a high degree of stability over the course of the session. B) Zoomed in portion of trace shown in (A). Note that the 405 nm channel shows negative deflections during positive deflections in the 465 nm channel, which is likely due to the fact that 405 nm differs from the isosbestic wavelength for GRAB-DA2m of 440 nm (Sun et al., 2020).

**Figure 7 - Figure Supplement 6:**
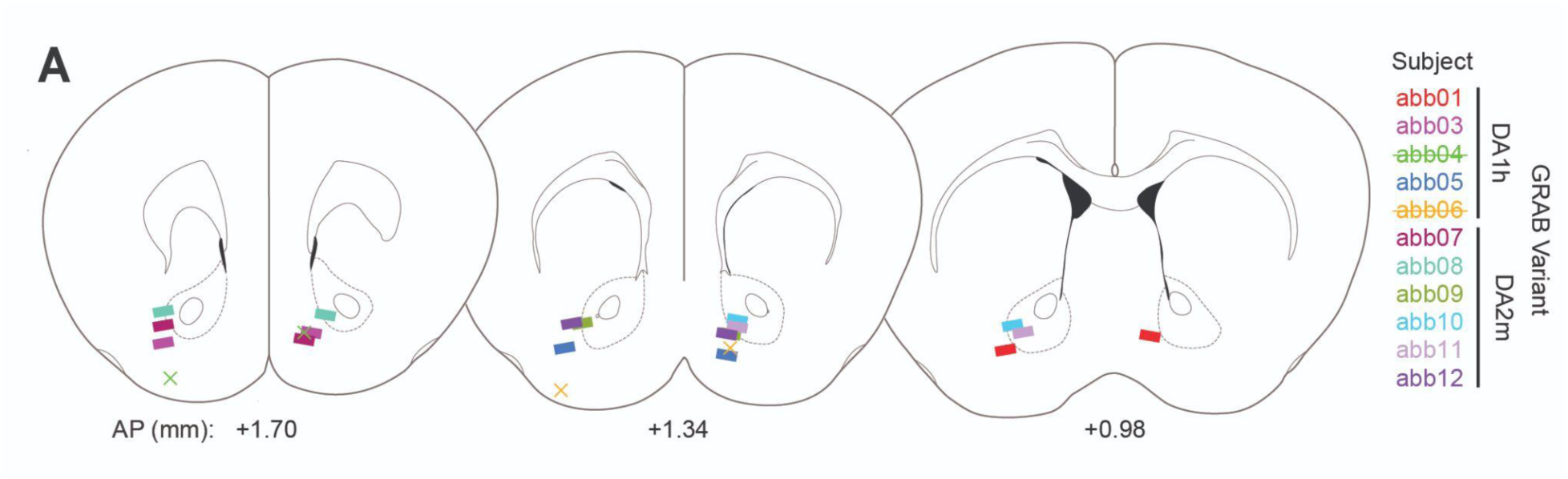
Fiber Placements for Fiber-Photometry. A) Position of fibers for fiber-photometry experiments shown in Figure 7 (AP relative to bregma). Rectangles depict the fiber position determined by histology for mice included in the analysis, while the “X” symbols depict the fiber position of two mice that were removed from the experiment for having missed NAcShL placements. In the experiment, the lateralization of placements for the NAcShL and NAcShM were randomized across mice. For this figure, fiber positions on the left side of the diagram depict fibers targeting in the NAcShL, and fibers on the right side of the diagram depict fibers targeting in the NAcShM.

## Notes

https://github.com/agordonfennell/OHRBETS

### Competing Interest Statement

The authors have declared no competing interest.

